# Chitinase-like proteins promoting tumorigenesis through disruption of cell polarity via enlarged endosomal vesicles

**DOI:** 10.1101/2022.08.17.504232

**Authors:** Dilan Khalili, Martin Kunc, Sarah Herbrich, Anna Mueller, Ulrich Theopold

## Abstract

Chitinase-like proteins (CLPs) are associated with tissue-remodeling and inflammation but also with several disorders, including fibrosis, atherosclerosis, allergies, and cancer. However, CLP’s role in tumors is far from clear. Here, we utilize *Drosophila melanogaster* to investigate the function of CLPs (imaginal disc growth factors; Idgf’s) in *Ras^V12^* dysplastic salivary glands. We find one of the Idgf’s members, *Idgf3*, is transcriptionally induced in a JNK-dependent manner via a positive feedback loop mediated by reactive oxygen species (ROS). Moreover, Idgf3 accumulates in enlarged endosomal vesicles (EnVs) that promote tumor progression by disrupting cytoskeletal organization. The process is mediated via the downstream component, αSpectrin, which localizes to the EnVs. Our data provide new insight into CLP function in tumors and identifies specific targets for tumor control.

## 1 Introduction

Chitinase-like protein (CLPs), including human YKL-39 and YKL-40 are synthesized and secreted under various conditions, including tissue injury, inflammatory and regenerative responses. Under pathological conditions they may contribute to asthma, sepsis, fibrosis and tumor progression (Roslind and Johansen 2009, Shao, Hamel et al. 2009) including ductal tumors, such as the lung, breast, and pancreas (Johansen, Jensen et al. 2006, Uhlen, Zhang et al. 2017). CLPs are regulated by growth factors, cytokines, stress and the extracellular matrix (ECM). However, the causal connection between CLPs’ function and disease progression is only partially elucidated (Park, Yun et al. 2020).

Animal models have been increasingly used in molecular oncology. This includes the fruitfly *Drosophila melanogaster*, where overexpression of dominant-active Ras (Ras^V12^) in proliferating tissue leads to benign tumors and simultaneous reduction of cell polarity genes to progression towards an invasive stage. (Brumby and Richardson 2003, Pagliarini and Xu 2003, Igaki, Pagliarini et al. 2006, Perez, Lindblad et al. 2017). Central to this switch towards increasing malignancy is the C-Jun N-terminal kinase (JNK)-signaling pathway, which becomes activated via loss of cell polarity and promotes tumor growth (Zhu, Xin et al. 2010). However, the outcome of activated JNK is mediated in a context-dependent manner due to downstream effects several of which are yet to be elucidated (Ciapponi, Jackson et al. 2001, Zeke, Misheva et al. 2016). Among potential JNK regulators, spectrin family members belong to cytoskeletal proteins which form a spectrin-based membrane skeleton (SBMS) (Bennett and Baines 2001). Through the Rac family of small GTPases, cell polarity and SBMS organization are maintained (Lee and Thomas 2011, Fletcher, Elbediwy et al. 2015). Although the exact relationship between Spectrin and JNK in tumors remains to be established, Rac1 under physiological conditions cooperates with JNK in tissue growth (Baek, Kwon et al. 2010, Wertheimer, Gutierrez-Uzquiza et al. 2012, Archibald, Mihai et al. 2015).

To explore CLPs’ tissue autonomous function in a ductal tumor, we utilize the *Drosophila melanogaster* salivary glands (SGs). Generally, *Drosophila* CLPs are endogenously expressed in the larvae and include six members, termed Idgf 1-6 (Imaginal disc growth factors), that are involved in development, establishment of the cuticle, wound healing and restoration of cell organization (Kirkpatrick, Matico et al. 1995, Kawamura, Shibata et al. 1999, Kucerova, Kubrak et al. 2016, Pesch, Riedel et al. 2016, Yadav and Eleftherianos 2018). The SGs’ epithelial luminal organization and the conserved activation of the tumor-promoting signaling factors make them suitable for dissecting CLP function. Moreover, the lumen separating a single layer of cells can be disrupted by constitutive active *Drosophila Ras* (*Ras^V12^*) (Krautz, Khalili et al. 2020) leading to the loss of ECM integrity, the formation of fibrotic lesions and of the oss of secretory activity (Khalili, Kalcher et al. 2021).

Here we investigated the role of *Drosophila* Idgf’s in *Ras^V12^*-expressing SGs. We show that one of the CLP’s members, *Idgf3*, is induced in tumor glands, leading to a partial loss of epithelial polarity and promoting a reduction of lumen size. The mechanism is driven through JNK signaling upstream of *Idgf3*. In line with previous work, ROS production via JNK mediates induction of *Idgf3*, creating a tumor-promoting signaling loop. Idgf3 further promotes the formation of enlarged endosomal vesicles (EnVs) via αSpectrin. Inhibiting EnVs formation by individually knocking-down *Idgf3* and *αSpectrin*, restores cell organization. Similar effects are observed upon expression of human CLP members in *Ras^V12^* SGs. Thus, our work identifies a phylogenetically conserved contribution of tumor-induced CLP’s towards the dysplasia of ductal organs and supports a role for spectrins as tumor modifiers.

## 2 Materials and Methods

### 2.1 *Drosophila* maintenance and larvae staining

Stocks were reared on standard potato meal supplemented with propionic acid and nipagin in a 25°C room with a 12 h light/dark cycle. Female virgins were collected for five days and crossed to the respective males (see supplementary cross-list) after two days. Eggs were collected for six hours and further incubated for 18 h at 29°C. 24 h after egg deposition (AED), larvae were transferred to a vial containing 3 mL food supplemented with antibiotics (see Table S1). 96 h and 120 h after egg deposition (AED), larvae were washed out with tap water before being dissected.

### 2.2 Sample preparation and immunohistochemistry

SGs were dissected in 1 x phosphate-buffered saline (PBS) and fixed in 4% paraformaldehyde (PFA) for 20 min. For extracellular protein staining, the samples were washed three times for 10 min in PBS and with PBST (1% TritonX-100) for intracellular proteins. Subsequently, samples stained for H2 were blocked with 0.1% bovine serum albumin (BSA) in PBS, and SG stained for pJNK, Idgf3, Spectrin, Dlg, p62 (ref(2)P), and GFP were blocked with 5% BSA for 20 min. After that, samples were incubated with the respective primary antibodies. Anti-pJNK (1:250), anti-Idgf3 (0.0134 μg/ml), anti-Spectrin (0.135 μg/ml) diluted in PBST were incubated overnight 4°C. anti-GFP (1 μg/ml) in PBST, H2 (1:5), and anti-SPARC (1:3000) in PBS were incubated for one hour at room temperature (RT). Samples were washed three times with PBS or PBST for 10 min and incubated with secondary antibody anti-mouse (4 μg/ml, Thermofisher #A11030) or anti-rabbit (4 μg/ml, Thermofisher #A21069) for one hour at RT. Subsequently, samples were washed three times in PBS or PBST for 10 min and mounted in FluoromountG.

### 2.3 Salivary gland size imaging and analysis

SG samples were imaged with Axioscope II (Objective 4x) (Zeiss, Germany) using AxioVision LE (Version 4.8.2.0). The images were exported as TIF and analyzed in FIJI (ImageJ: Version 1.53j). Representative confocal pictures were selected for figure panels and the complete set of replicate figures processed further for quantification (see below). Region of Interest (ROI) were drawn with the Polygon selection tool, and the scale was set to pixels (Px). The SG area was summarized as a boxplot with whisker length min to max. The bar represents the median. Statistical analysis was done with Prism software (GraphPad Software, 9.1.2, USA), the population was analyzed for normality with D’Agostino-Pearson and p-value quantified with Student’s t-test.

### 2.4 Nuclear volume imaging and quantification

Nuclei were stained with DAPI (1 μg/ml, Sigma-Aldrich D9542) in PBST for 1 h at RT. Mounted glands were imaged with Zeiss LSM780 (Zeiss, Germany) using a plan-apochromat 10x/0.45 objective with a pixel dwell 3.15 μs and 27 μm pinhole in z-stack and tile scan mode. Zeiss images were imported into ImageJ and viewed in Hyperstack. The selection threshold was set individually for each sample, and the analysis was performed with 3D objects counter. The nuclei volume was presented in boxplot, whisker length min to max and bar represent median. P-value quantified with Student’s t-test and the scale bar represent μm^3^.

### 2.5 Intensity and hemocyte quantification

The images for quantifying pJNK, TRE, Idgf3, and SPARC intensity and hemocyte recruitment were captured with AxioscopeII (Objective 4x) (Zeis, Germany). The images were exported as TIF and analyzed in ImageJ. ROI was drawn with the Polygon selection tool, and subsequently, the total intensity was measured (pixel scale). The intensity was quantified according to the equation: Integrated Density – (SG area*Mean gray value). Hemocyte area was selected with Threshold Color and quantified by using the following equation: Ln (Hemocyte area + 1)/Ln (SG size + 1). Representative images were taken with Zeiss LSM780 (Zeiss, Germany). The images were then processed using Affinity Designer (Serif, United Kingdom). Graphs and statistical analysis were generated with Prism software (GraphPad Software, 9.1.2, USA). The population was analyzed for normality with D’Agostino-Pearson. Statistical significance was determined with Student’s t-test, One-way ANOVA with Tukey’s multiple comparison, and two-way ANOVA with Dunnett’s multiple comparison.

### 2.6 Enlarged endosomal vesicles penetrance quantification

The penetrance of the enlarged vesicles was subjectively quantified. Samples were analyzed in Axioscope II (Objective 20x) (Zeiss, Germany). At least 15 samples were analyzed with three independent replicas.

### 2.7 Humanized transgenic *Drosophila* lines

Plasmids were generated and transformed at VectorBuilder (https://en.vectorbuilder.com/). Human *CH3L1* and *CH3L2* genes were inserted into *Drosophila Gene Expression Vector pUASTattB* vector generating VB200527-1248haw and VB200518-1121xyy, respectively and transformed into *E. coli*. The bacteria were cultured in 3 ml LB supplemented with ampicillin (AMP: 100 ug/ml) for 15 h, at 37°C. The plasmid was extracted according to the GeneJetTm Plasmid Miniprep Kit #K0503 standard procedure. Plasmids were validated through sequencing at eurofins (https://www.eurofins.se/: For primer details see Table S1). *Drosophila* transgenic lines were generated at thebestgene (https://www.thebestgene.com/). Plasmids were extracted with QIAGEN Plasmid Maxi Kit according to the standard procedure and injected into *w^1118^* strains. Expression of the human CLPs was validated with qPCR.

### 2.8 *In situ* hybridization

The Idgf3 (GH07453: DGRC) probe was generating according to (Hauptmann., 2015). The staining procedure is described elsewhere with the following changes (Hauptmann et al., 2016). The procedure was conducted in 200 μl transwells containing four salivary glands. The procedure included three technical replicas per genotype. Images were aqured with Leica MZ16 (Leica, Germany) microscope and Leica DFC300x FX digital color camera (Leica, Germany). Representative images were taken, and figures were generated in Affinity Designer (Serif, United Kingdom).

### 2.9 qPCR

mRNA isolation and cDNA synthesis were performed according to manufacture instructions (AM1931). qPCR procedures were performed as described earlier (Krautz et al., 2021) with an adjusted Kappa concentration to 0.5x. At least three replicates and two technical replicates were performed for each qPCR. See supplementary Table S1 for primer list.

## 3 Results

### 3.1 Idgf3 promotes a dysplastic phenotype

Obstruction of SG lumen by the constitutive-active oncogene, *Ras^V12^*, under *Beadex-Gal4* driver (*Ras^V12^*) disrupts organ function between 96 h and 120 h after egg desposition (AED) (Khalili, Kalcher et al. 2021). Being that CLPs have been implicated in the loss of cell polarity (Morera, Steinhauser et al. 2019), we investigated whether *Drosophila* CLPs contribute to the observed phenotype. First, to find out whether CLPs were induced in the *Ras^V12^* glands, we assessed relative mRNA levels at two different time points, 96 h and 120 h AED. Only one of the *CLP* members, namely *Idgf3*, was significantly upregulated at both time points (Fig. 1A and Fig. S1A). Therefore, we decided to focus on *Idgf3’s* effects on dysplastic glands.

**Figure 1.**
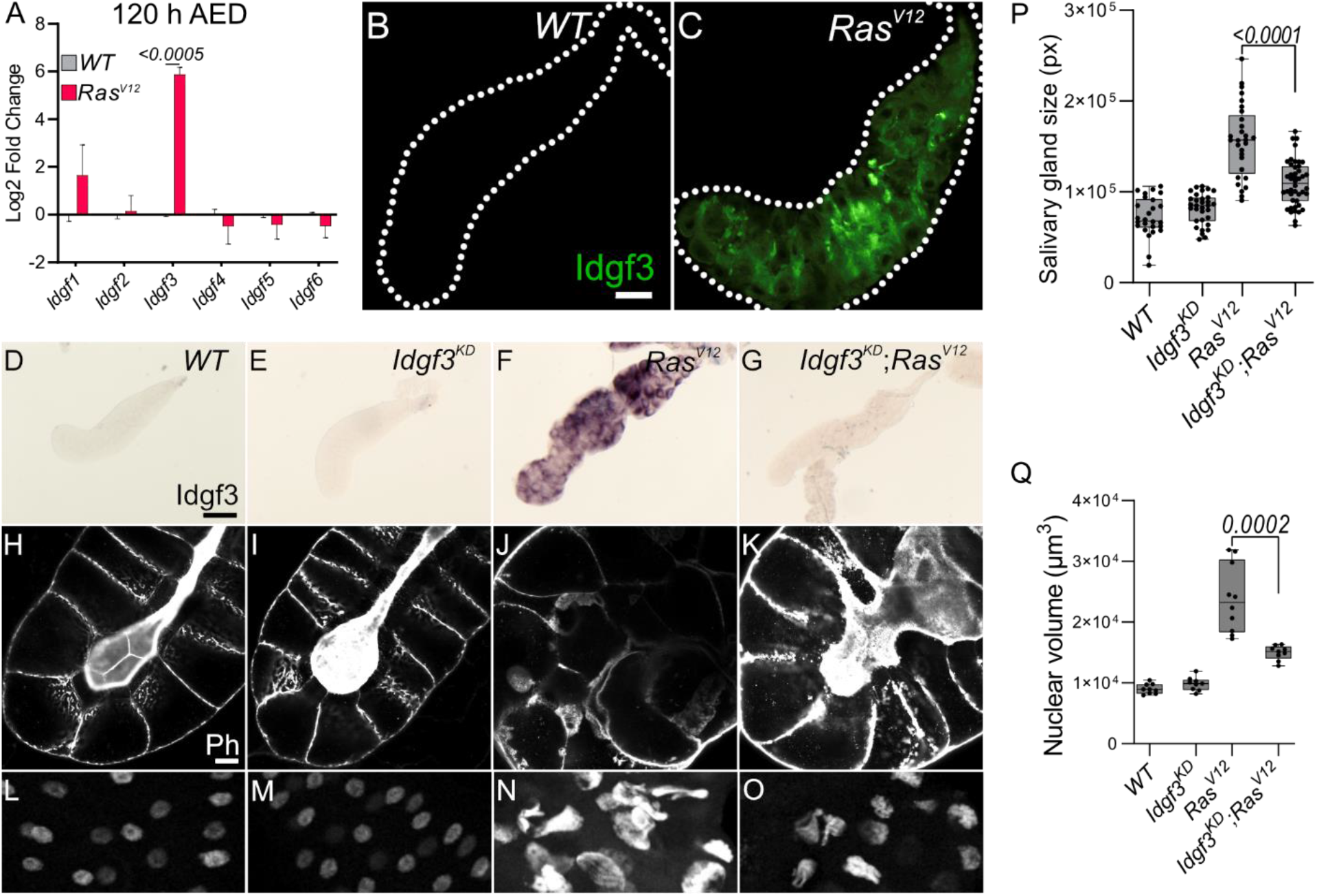
Idgf3 promotes growth and disrupts tissue architecture. **(A)** qPCR data showing induction of *Idgf3* in 120 h AED *Ras^V12^* glands. **(B-C)** Idgf3 tagged with GFP was localized in the dysplastic glands. **(D-G)** Knock-down of *Idgf3* in *Ras^V12^* glands confirmed reduced mRNA levels as shown by in situ hybridization. **(H-K)** F-actin (Phalloidin) staining revealed partial restoration of the lumen in *Idgf3^KD^;Ras^V12^* glands, in comparison to *Ras^V12^* alone. **(P)** SG size quantification showing a reduction in tissue size in *Idgf3^KD^;Ras^V12^* SG compared to *Ras^V12^* alone. **(L-O)** Nuclei in DAPI stained SG displayed a reduced size in *Idgf3^KD^;Ras^V12^;* quantified in **(Q)**. Scale bars in **(B-C)** represent 100 μm, **(D-G)** represent 0.3 mm and **(H-K)** represent 20 μm. Data in **(A)** represent 3 independent replicas summarized as mean ± SD. Boxplot in **(P, Q)** represent at least 20 SG pairs. Whisker length min to max, bar represent median. P-value quantified with Student’s t-test.

Idgf3 contains an N-terminal signal peptide and has been detected in hemolymph (Karlsson, Korayem et al. 2004). To analyze its subcellular tissue distribution in SGs, we used a C-terminally GFP tagged version of Idgf3 (Kucerova, Kubrak et al. 2016). At 96 h we could not detect Idgf3 in the whole *WT* or *Ras^V12^* animals (Fig. S1F-G’), possibly due to limited sensitivity. Likewise, 120 h old *WT* larvae did not show any detectable signal (Fig. S1H-H’) while a strong Idgf3 signal was detected in *Ras^V12^* SGs (Fig. S1I-I’). To better understand Idgf3 distribution at a higher resolution, we dissected 120 h AED glands. *WT* glands had a weaker Idgf3∷GFP signal in comparison to the *Ras^V12^* (Fig. 1B-C). Moreover, Idgf3 was unevenly distributed throughout *Ras^V12^* SGs (Fig. 1C).

The increased level of *Idgf3* between 96 h and 120 h strongly correlated with loss of tissue- and cell-organization and an increased nuclear volume (Krautz, Khalili et al. 2020). In order to characterize the role of Idgf3 in *Ras^V12^* glands, we used a specific *Idgf3 RNA-interference* line (*Idgf^KD^*). Moreover, we focused on 120 h larvae, unless otherwise stated, since they showed the most robust and developed dysplastic phenotype. Efficient knockdown of *Idgf3* was confirmed using ISH and at the protein level (Fig. 1D-G, S1J-M; quantified in N, (Kucerova, Kubrak et al. 2016)). Macroscopic inspection showed that *Idgf^KD^;Ras^V12^* SGs were smaller than *Ras^V12^* SGs (Fig. 1P), resembling WT controls. To gain insight into the cellular organization, we stained the glands for F-actin (Phalloidin: Ph) and DNA using DAPI. In *Idgf^KD^* the cells retained their cuboidal structure, and the lumen was visible as in *WT*, indicating that Idgf3 on its own does not affect apicobasal polarity (Fig. 1H-I). In contrast, in *Ras^V12^* glands apicobasal polarity was lost, and the lumen was absent (Fig. 1J, (Khalili, Kalcher et al. 2021). In *Idgf^KD^;Ras^V12^* SGs a reversal to the normal distribution of F-actin and partial restoration of the lumen was observed (Fig. 1K). Similarly, the nuclear volume, which increased in *Ras^V12^* SGs returned to near wild type levels upon *Idgf^KD^* (Fig. 1 L-O, quantified in Q). This indicates that *Idgf^KD^* can rescue Ras^V12^-induced dysplasia.

In order to unravel the specific effects mediated by Idgf3 we further investigated *Ras^V12^* associated phenotypes, including fibrosis and the cellular immune response. As recently reported, *Ras^V12^* SGs displayed increased levels of the extracellular matrix components (ECM), including collagen IV and SPARC (BM40, (Khalili, Kalcher et al. 2021)). *Idgf^KD^* did not affect SPARC levels in comparison to the *WT* (Fig. S1 O-P) but *Idgf-KD;Ras^V12^* SGs displayed significantly reduced SPARC levels in comparison to *Ras^V12^* (Fig. S1 Q-R, quantified in S). To assess whether this led to a reduced inflammatory response, we investigated the recruitment of plasmatocytes, macrophage-like cells previously reported to be recruited towards tumors (Perez, Lindblad et al. 2017). We found that both control and *Idgf^KD^* glands did not show recruitment of hemocytes (Fig. S1T-U). In contrast to the effects on ECM components, *Idgf^KD^ in Ras^V12^* glands did not lead to any changes in hemocyte attachment (Fig. S1V-W, quantified in X). Taken together, upon *Ras^V12^* overexpression, Idgf3 promotes SG overgrowth, loss of cell organization, and fibrotic-like accumulation of the ECM, but not immune cell recruitment.

### 3.2 Idgf3 induces dysplasia via JNK-signaling

Dysplasia is driven by internal and external factors that either work in concert or independently. Similar to what we observed in *Idgf^KD^;Ras^V12^* glands blocking the sole *Drosophila* JNK member *basket* reverts many tumor phenotypes. Moreover, the dysplastic loss of apical and basolateral polarity between 96 h and 120 h is driven by the JNK-pathway (Krautz, Khalili et al. 2020). The time frame when we observed upregulation of *Idgf3* (Fig. 1A, S1A) coincides with the period during which blocking JNK restores tissue organization and homeostasis, similar to what occurs in *Idgf^KD^;Ras^V12^* tissues (Fig. 1K, S1R). Therefore, we decided to test a possible involvement of JNK-signaling in the regulation of Idgf3.

First, we performed a targeted JNK RNAi-screen using Idgf3∷GFP intensity in the glands as readout upon KD of JNK signaling components. We first confirmed the sensitivity of the Idgf3∷GFP construct by *Idgf3-KD* in *Ras^V12^* SGs compared to *Ras^V12^* glands (Fig. S2A-B, quantified in Fig S2C). KD of the two classical TNF receptors upstream of JNK, *Grnd* (*Grindelwald*) and *Wgn* (*Wengen*) similarly reduced Idgf3∷GFP intensity (Fig 2B-C, quantified in E (Palmerini, Monzani et al. 2021)). Similar effects were observed with *Bsk^KD^* (Fig. 2A, D, quantified in E). Altogether this suggests that Idgf3 protein levels are regulated downstream of JNK and the TNF members *Grnd* and *Wgn*.

**Figure 2.**
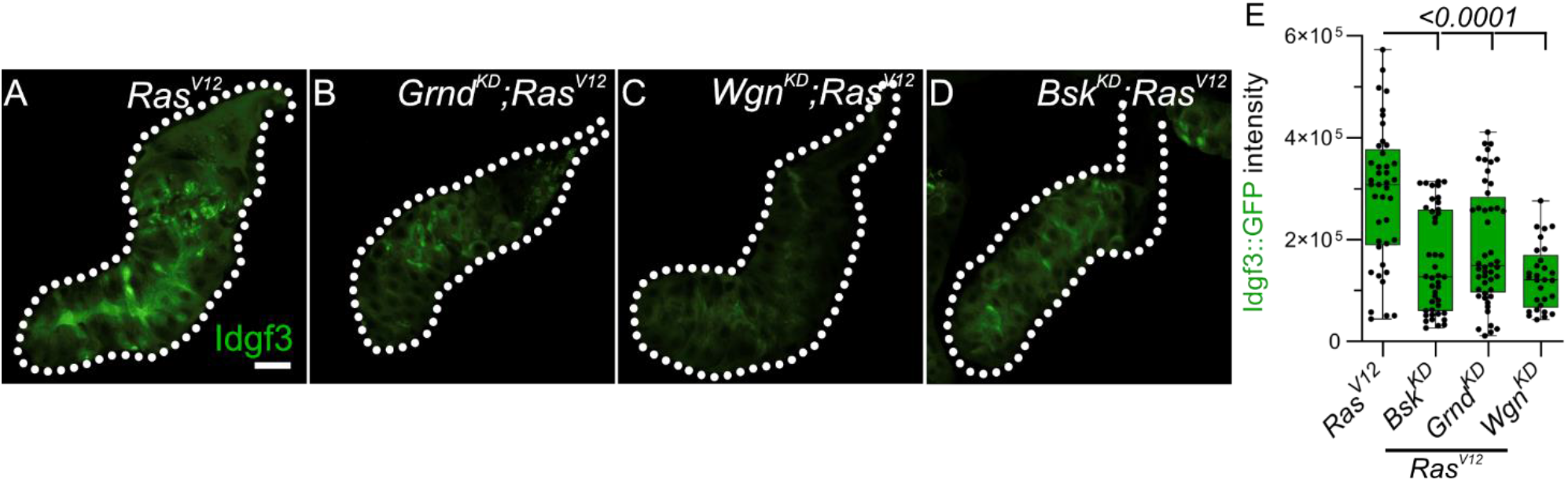
Idgf3 dysplasia is mediated through JNK activity. **(A-D)** Representative images of Idgf3∷GFP in a JNK targeted screen. **(E)** Quantification showing Idgf3∷GFP intensity was reduced by *Grnd^KD^*, *Wgn^KD^* and *Bsk^KD^* in *Ras^V12^* SG. Scale bars in **(A-D)** represent 100 μm. Boxplot in **(E)** represents at least 20 SG pairs. Whisker length min to max, bar represent median. P-value quantified with Student’s t-test.

### 3.3 ROS promotes Idgf3 induction via JNK

To further dissect Idgf3 regulation, we focused on the positive JNK regulators, reactive oxygen species (ROS) both intra- and extracellularly (Diwanji and Bergmann 2017, Perez, Lindblad et al. 2017). We previously reported that ROS production in *Ras^V12^* SGs increases via JNK (Krautz, Khalili et al. 2020). To inhibit ROS intra- and extracellularly, we separately overexpressed the H_2_O_2_ scavengers Catalase (Cat) and a secreted form of Catalase, IRC (immune-regulated Catalase), and O_2_-scavenger SOD (Superoxide dismutase A), in the *Ras^V12^* background and quantified Idgf3∷GFP intensity. Reducing levels of intracellular H_2_O_2_ (*Cat^OE^*), but not O_2_-(*SOD^OE^*) lowered Idgf3∷GFP intensity (Fig. S3A-D quantified in E). Similarly, reduction of extracellular H_2_O_2_ by the secreted version of Catalase lowered Idgf3∷GFP levels (Fig. 3A-D, quantified in E) as well as JNK signaling (Fi. 3J, K-N’ quantified in O). In line with the reduced tissue size and improved tissue integrity in *Idgf3^KD^;;Ras^V12^*, overexpression of *IRC* in *Ras^V12^* SGs also reduced SG size (Fig. 3T), improved tissue integrity and restored the SG lumen (Fig. 3F-I, P-S).

**Figure 3.**
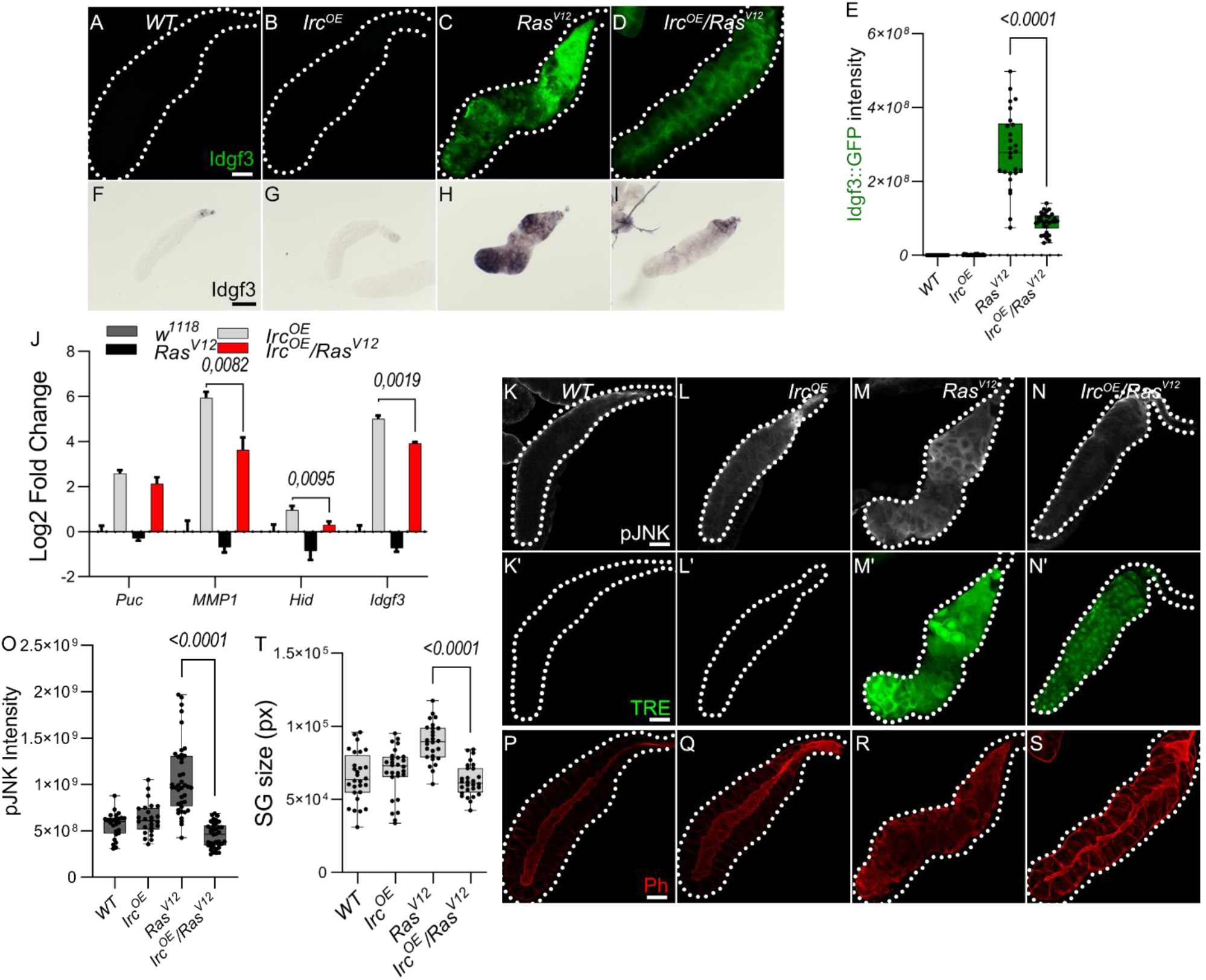
Idgf3 regulation feeds into a JNK-ROS feedback loop. **(A-D)** Reduction of H_2_O_2_ by overexpression of secreted catalase (immune regulated catalase; IRC) lowered Idgf3∷GFP levels, quantified in **(E)**. **(F-I)** ISH showing reduced expression of *Idgf3* in *IRC-OE;Ras^V12^*glands. **(J)** qPCR data showing reduction of *Idgf3, MMP1* and *Hid* in *IRC^OE^;Ras^V12^*glands. **(K-N’)** pJNK staining and TRE reporter constructs showing reduced intensity in *IRC^OE^;Ras^V12^* in comparison to *Ras^V12^* glands, quantified in **(O)**. **(P-S)** Phalloidin staining showing partially restored lumen in *IRC^OE^;Ras^V12^* glands, quantified in **(T)**. Scale bars in **(A-D, K-S)** represent 100 μm and **(F-I)** represent 0.3 mm. Data in **(J)** represent 3 independent replicas summarized as mean ± SD. Boxplot in **(E, O, T)** represent at least 20 SG pairs. Whisker length min to max, bar represent median. P-value quantified with Student’s t-test.

In summary, ROSs contribute to pJNK signaling. In addition, overexpression of extracellular and intracellular Catalase but not SOD reduces Idgf3 induction via JNK, similar to the feedback loop that has been identified in other tumor models (Perez, Lindblad et al. 2017).

### 3.4 Idgf3 accumulates in large vesicles, which display markers for endocytosis and macropinocytosis

We previously noted the uneven distribution of Idgf3 in *Ras^V12^* SGs (Fig. 1C). To further understand how Idgf3 promotes dysplasia, we dissected its subcellular localization (Fig. 4A). We stained the glands for F-actin (Phalloidin) and addressed Idgf3∷GFP localization at high resolution (Fig. 4B-C’). Interestingly, we observed Idgf3∷GFP clusters surrounded by F-actin (Fig. 4C-C’: arrow). Using a different salivary gland driver (*AB-Gal4*) to drive expression of *Ras^V12^*, we also observed increased expression of Idgf3∷GFP and its localization within vesicular structures (Fig. S4V-W’: arrow). The size of the vesicle-like structures was between 10-43 μm in comparison to secretory *Drosophila* vesicles (3-8μm, Fig. 4D) (Tran and Hagen., 2017). We refer to these as enlarged vesicles (EnVs). Based on the increased Idgf3 levels, we wondered whether the protein was aggregating in EnVs. Unlike in *WT* glands, the aggregation marker, p62 (*Drosophila* Ref(2)P, (Bartlett, Isakson et al. 2011) strongly bound to the cytoplasm of *Ras^V12^ SGs*. However, the EnVs did not contain any aggregated proteins (Fig. S4X-Y’).

**Figure 4.**
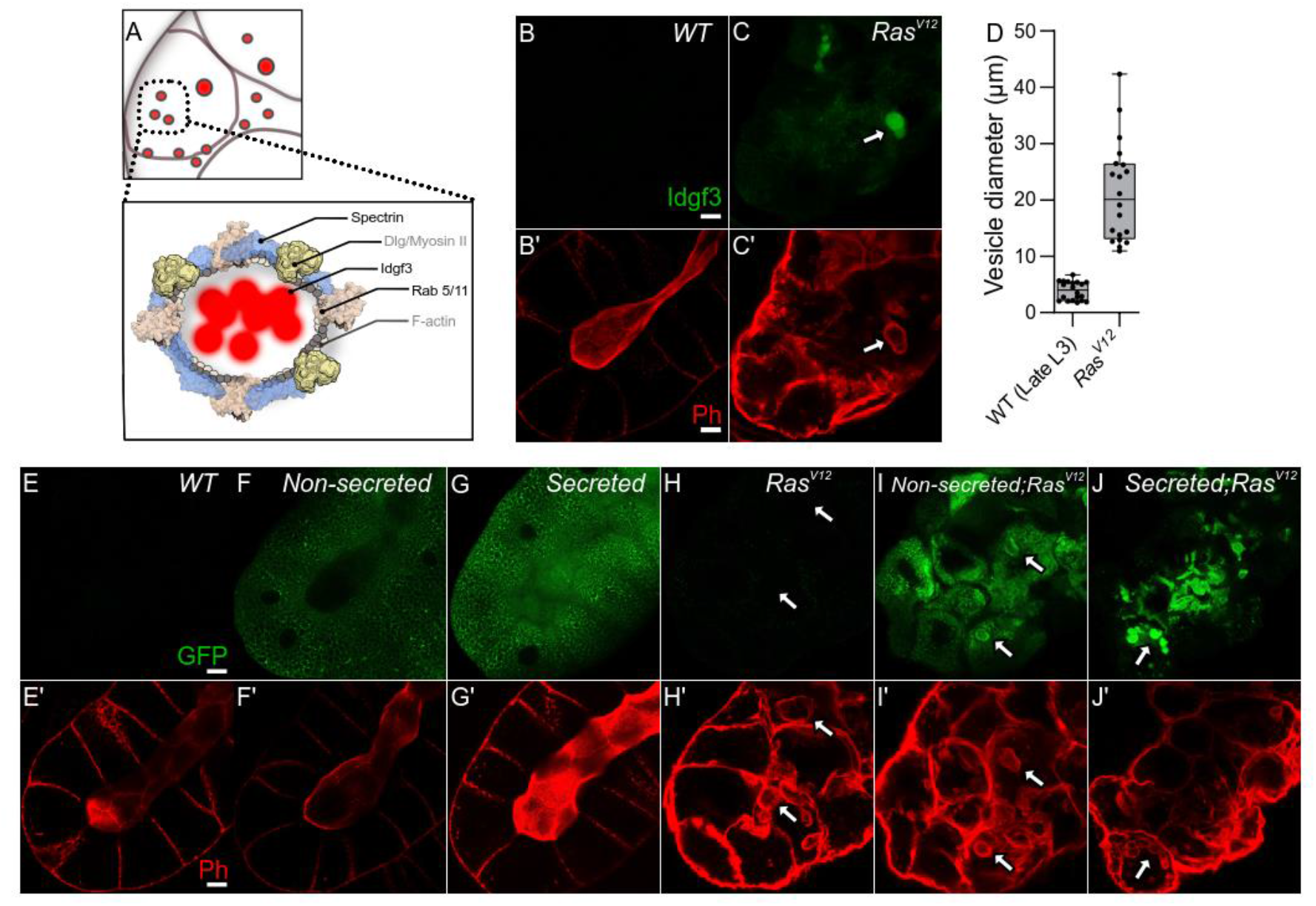
*Idgf3* promotes formation of enlarged endosomes. **(A)** Idgf3 enclosed by enlarged vesicles (EnVs) coated by cytoskeletal and cell polarity proteins. **(B-C’)** Idgf3∷GFP clusters coated with Phalloidin. **(D)** Vesicle size quantification showing *Ras^V12^* enlarged vesicles in comparison to prepupae SG vesicles. **(E-J’)** Non secreted MFGE8 localizes to the surface of EnVs, co-stained with phalloidin. The secreted MFGE8 is packaged into EnVs in *Ras^V12^*glands. Scale bars in **(B-C’, E-J‘)** represent 20 μm. Boxplot in **(D)** represents 20 EnVs. Whisker length min to max, bar represent median. P-value quantified with Student’s t-test.

To further understand whether the localization of Idgf3 in EnVs was dependent on the presence of a secretion signal we overexpressed two versions of human phosphatidylserine binding protein, MFG-E8 (Milk fat globule-EGF factor), without (referred as non-secreted: Fig .4F-F’,I-I’) and with a signal peptide (referred as secreted: Fig. 4G-G’,J-J’, Asano et al., 2004). In controls, the non-secreted form was found in the cytoplasm, whereas the secreted version was detected in the cytoplasm and in the lumen (Fig. 4F-G’). In *Ras^V12^* SGs, the non-secreted form was surrounding the EnVs (arrow), indicating the presence of phosphatidylserine on their membrane (Fig. 4I-I’). In contrast, the secreted form localized to the inside of the EnVs (Fig. 4J-J’: arrow). These data suggest that EnVs are surrounded by a lipid membrane and recruit from the SG lumen.

In order to further characterize Idgf3-containing EnVs we co-expressed vesicle-specific Rab’s coupled with a GFP fluorophore, a lysosomal marker (Atg8), an autophagy marker (Vps35), and a marker for phosphatidylinositol-3-phosphate-(PtdIns3P: *FYVE*)-positive endosomes in *Ras^V12^* glands (For a complete set, see Fig. S4A-I’’). To increase sensitivity and to identify EnVs, we stained with anti-GFP and co-stained with Phalloidin. Localization of Rabs and phalloidin to the same vesicles was observed with Rab5 and Rab11 but not Rab7 (Fig. S4A-D’’, S4I-I’’). Moreover, EnVs were also positive for PtdIns3 (Fig. S4H-H’’). In line with their dependence on secretion, this potentially identifies EnVs as enlarged recycling endosomes. EnV accumulation in *Ras^V12^* glands between 96 h and 120 h implies that (i) endosome formation is either increased compared to *WT* or (ii) that endosomes are not normally recycled leading to their accumulation. The latter hypothesis correlates with the loss of apico-basolateral polarity and the disruption of secretion due to a lack of a luminal structure (Khalili, Kalcher et al. 2021). To test the first hypothesis, we blocked the formation of early endosomes with *Rab5^DN^*. Apico-basolateral polarity, detected by a visible lumen, was not affected by *Rab5^DN^*. Moreover, *Rab5^DN^;Ras^V12^* did not block EnV formation and restoration of apicobasal polarity (Fig. S4J-M). Halting the recycling endosome pathway via *Rab11^DN^* increases the endosomes’ accumulation without affecting cell polarity (Fig. S4N). In contrast, in *Rab11^DN^;Ras^V12^* SGs, endosomes were not accumulating, and EnVs were still detected (Fig. S4O). Taken together, EnV formation is independent of the classical recycling pathway, suggesting other candidates are involved in their generation.

In SGs, overexpression of Rac generates enlarged vesicles with similarity to the EnVs described here (Lee and Thomas 2011). Supporting a role in dysplasia in our system, *Ras^V12^* SGs showed stronger Rac1 expression in comparison to the control. Moreover, we observed Rac1 also localized to EnVs (Fig. S4P-Q’). Decoration with Rac1 and actin as well as their dependence on Ras activation potentially identifies EnVs as macropinocytotic vesicles ((Recouvreux and Commisso 2017), see also discussion). The enlarged vesicles that form upon Rac overexpression in SGs (Lee and Thomas 2011) also stain positive for Spectrins identifying them as additional candidates for EnVs formation. Of note, Spectrins under physiological settings are involved in the maintenance of cellular integrity including epithelial organization, which is lost in *Ras^V12^* SGs.

### 3.5 JNK promotes EnVs formation via Idgf3 upstream of αSpectrin

To analyze Spectrin contribution to EnVs formation, we stained for αSpectrin, one of the three members in flies (Williams, Smith et al. 2003) and found it to be induced in *Ras^V12^* SGs and to localize to the EnVs (Fig. S5A-B’’). Kockdown of *Idfg3* in *Ras^V12^* SGs reduced both αSpectrin levels and EnVs formation (Fig 5A-D’). Despite efficient *Idfg3^KD^*, transcript levels for both α- and β_Heavy_Spectrin as well as for Rac1 were not affected indicating regulation at the posttranscriptional level (Fig. 5E). Moreover, we found markers for cell polarity including Dlg, and Myosin II also decorate the EnVs (Fig. S4R—U’: arrow). In contrast, αSpectrin^KD^ (Fig. S5C-F quantified G) reduced Idgf3 levels (Fig. 5F-I’ quantified J) as well as JNK signaling upstream of Idgf3 (Fig 5U-V). Further supporting a role for Spectrins in SG dysplasia, k/d of αSpectrin in *Ras^V12^* glands abolished EnVs formation and partially restored the SG lumen (Fig. 5I’’’).

**Figure 5.**
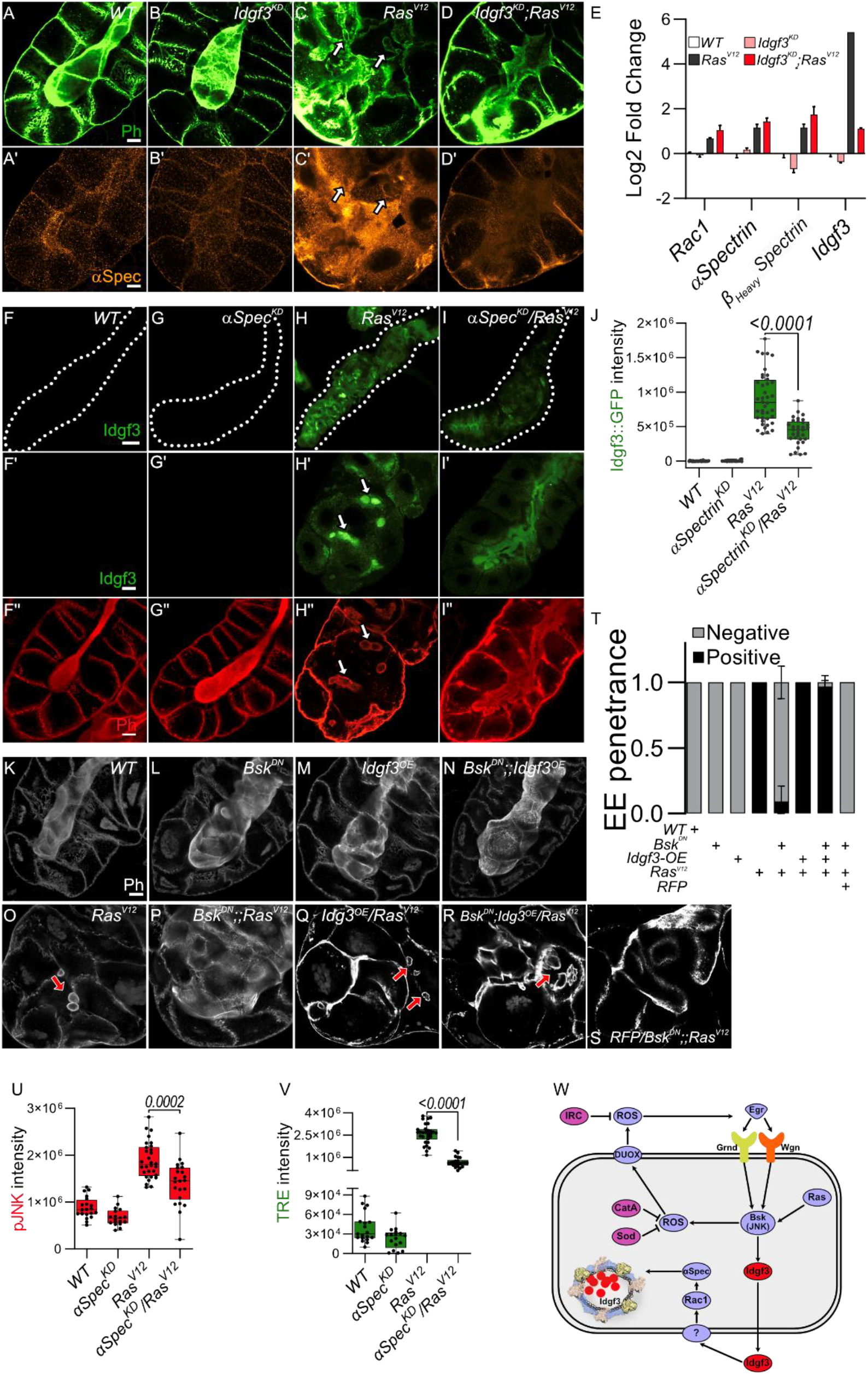
JNK promotes EnVs formation via Idgf3 upstream of αSpectrin. **(A-D’)**αSpectrin staining showing restoration of normal distribution in *Idgf3^KD^;Ras^V12^*glands. **(E)** qPCR data showing no reduction of *αSpectrin* and *β_Heavy_Spectrin* in *Idgf3^KD^;Ras^V12^*. **(F-I‘‘)** Reduced levels of αSpectrin (αSpectrin^*KD*^ /*Ras^V12^*) reduces Idgf3∷GFP levels quantified in **(J)**, prevents formation of EnVs and largely restores the SG lumen (arrows indicate EnVs). **(K-S)** Phalloidin staining showing epistasis of EnVs formation in which Idgf3 acts downstream of JNK. **(T)** EnVs penetrance quantification showing a strong induction of EnVs in *JNK^DN^;;Idgf3^OE^/Ras^V12^* glands. **(U)** pJNK intensity quantification showing reduced levels in *aSpectrin^KD^ /Ras^V12^*. **(V)** TRE intensity quantification showing reduced levels in *aSpectrin^KD^ /Ras^V12^*. **(W)** Idgf3 promotes formation of EnVs, upstream of Rac1. Scale bars in **(A-D’, F‘-I‘‘, K-S’)** represent 20 μm, **(F-I)** represents 100 μm. Data in **(E)** represent 3 independent replicas summarized as mean ± SD. Barplot in **(T)** represent 3 independent replicas with at least 10 SG pairs, summarized as mean ± SD. Boxplot in **(J, U-V)** represent at least 20 SG pairs. Whisker length min to max, bar represent median. P-value quantified with Student’s t-test.

Taken together this suggests that Idgf3 promotes EnVs formation (Fig. 5C-D) most likely post-transcriptionally (Fig. 5E). In line, overexpression of Idgf3 throughout the whole gland, at 96 h, as shown by ISH (Fig. S5I-L), led to an increase in the number of glands with endosomes (Fig. S5M-P’’’, quantified in Q). To address epistasis between Idgf3 and JNK we calculated the penetrance of EnVs formation. In *Ras^V12^* SGs we observed EnVs in 100 % of the glands, an effect that was strongly blocked in *Bsk^DN^;;Ras^V12^* (Fig. 5O-P, quantified in T). Blocking JNK and overexpressing *Idgf3* in *Ras^V12^* strongly reverted the *Bsk^DN^;;Ras^V12^* phenotype, a lumen could not be detected, and around 98% of the glands contained enlarged endosomes (Fig. 5O,R, quantified in T) while control SGs using RFP-overexpression retained the *Bsk^DN^;;Ras^V12^* phenotype. Overexpression of Idgf3 alone did not result in EnVs formation (Fig. 5K-N). In conclusion, the data suggest that Idgf3 acts downstream of JNK and - through formation of EnV’s - disrupts luminal integrity.

### 3.6 Human CLP members enhance dysplasia in *Drosophila* SGs

Finally, we wished to determine whether the tumor-modulating effects we had observed for *Drosophila* Idgf3 also applies to human CLP members. For this we expressed two human *CLPs* (*Ch3L1* or *Ykl-40*; 29% AA identity to Idgf3 and *Ch3L2* or *Ykl-39*; 26% AA identity, Fig. 6A) in SGs, both on their own and in combination with *Ras^V12^*. Similar to Idgf3, both CLPs enhanced the hypertrophy observed in *Ras ^V12^* SGs (Fig. 6B-G quantified in N). Additionally, *Ch3L1* enhanced the prevalence of EnVs in the Ras mutant background (Fig. 6O). Taken together this means that the tumor-promoting effect of CLPs is conserved between *Drosophila* and humans and may affect different phenotypes of dysplasia depending on the CLP under study.

**Figure 6.**
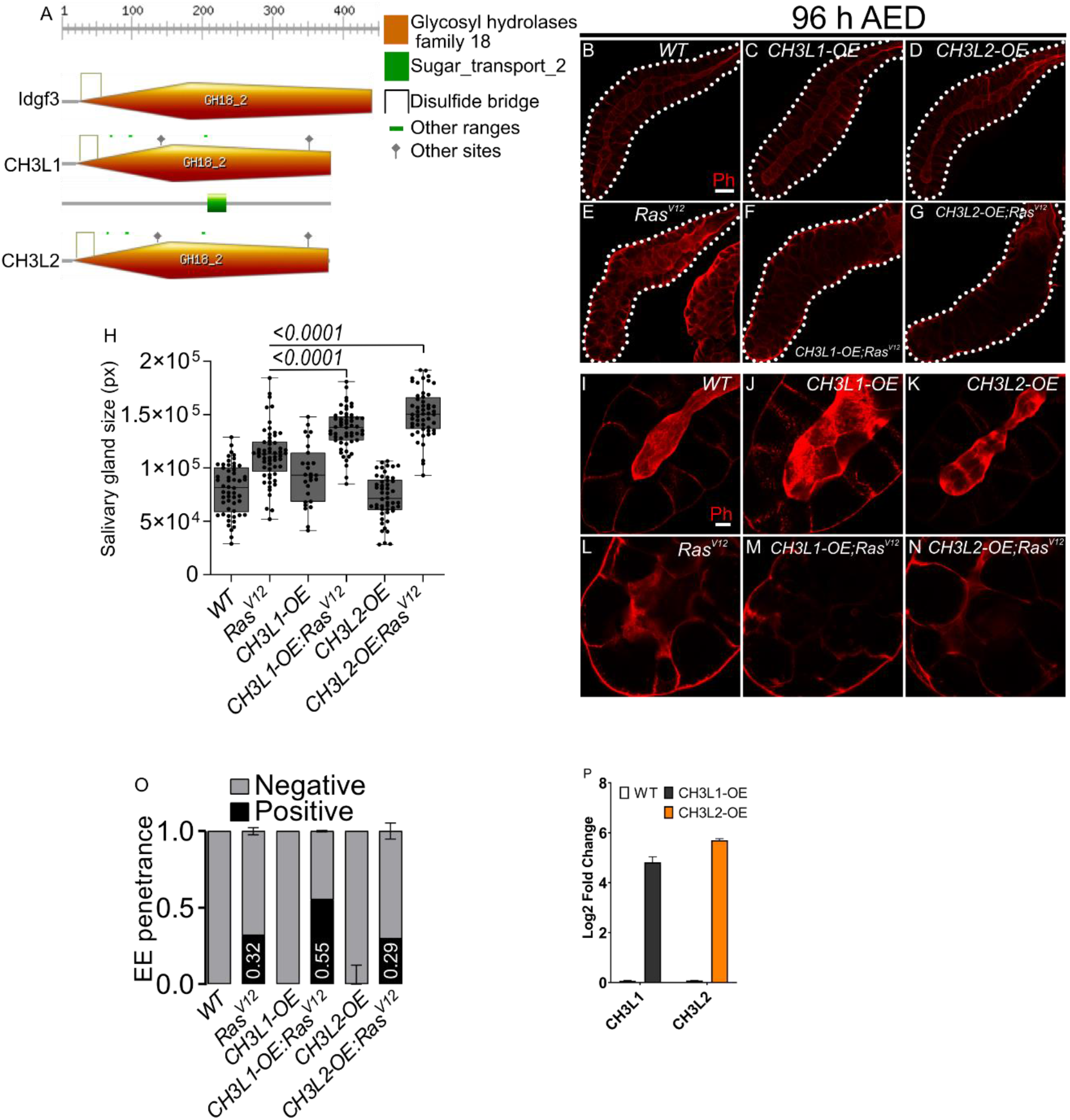
Human Chitinase-like proteins similarly to Idgf3 promotes EnVs formation. **(A)** Comparison of Idgf3, CH3L1 and CH3L2 protein motifs (https://prosite.expasy.org). **(B-G)** Representative images of phalloidin staining used for size quantification. **(H)** SG size quantification showing an increase in tissue size in *CH3L1^OE^;Ras^V12^* and *CH3L2^OE^;Ras^V12^* SG compared to *Ras^V12^* alone. **(I-N)** Phalloidin staining depicting disrupted lumen integrity in *Ras^V12^* glands. **(O)** EnVs penetrance quantification showing an induction of EnVs in *CH3L1^OE^;Ras^V12^* glands. **(P)** qPCR confirmation of CH3L1 and CH3L2 expression in SG. Scale bars in **(B-G)** represent 100 μm and **(I-N)** 20 μm. Boxplot in **(H)** represent at least 20 SG pairs. Whisker length min to max, bar represent median. P-value quantified with Student’s t-test. Barplot in **(O)** represent 3 independent replicas with at least 10 SG pairs, summarized as mean ± SD. Barplot in **(P)** represent 4 independent replicas with at least 10 SG pairs, summarized as mean ± SD.

## 4 Discussion

The levels of Chitinase-like proteins (CLPs) are elevated during a wide range of inflammatory processes as well as neoplastic disorders. Their physiological function has been more elusive but includes the formation of extracellular assemblages (Zhao, Su et al. 2020) including the insect cuticle (Pesch, Riedel et al. 2016), wound healing and in both mammals (Zhao, Su et al. 2020) and insects (Kucerova, Broz et al. 2015) and the restoration of cell integrity after oxidative damage (Lee, Da Silva et al. 2011). Conversely, induction of CLPs has been associated with the development of fibrotic lesions and cancer development with poor prognosis (reviewed in (Zhao, Su et al. 2020)). We used *Drosophila* as a tumor model to dissect CLP (*Idgf3*) function genetically in a secretory ductal organ, the salivary glands. We show that Idgf3 promotes tumor overgrowth through the disruption of cell polarity. The induction of *Idgf3* disrupts cell organization and leads to the formation of enlarged endosome vesicles (EnVs) which accumulate in the cytoplasm. Genetically, *Idgf3* is induced via a pro-tumorigenic JNK and ROS signaling feedback loop. Consequently, Idgf3 recruits the spectrin-based membrane skeleton (SBMS) for the formation of EnVs. Significantly, KD of *Idgf3* inhibits overgrowth, restores cell polarity, reduces ECM size and blocks EnVs formation.

Our identification of a contribution of JNK signaling and both extra- and intracellular ROS to dysplasia is in line with previous findings from other *Drosophila* tumor models (Fogarty and Bergmann 2017). Similarly, like others (Fogarty and Bergmann 2017) we observe an amplification loop between ROS and JNK signaling, which augments the dysplastic phenotype ((Krautz, Khalili et al. 2020) and this work). Several studies have demonstrated that activation of JNK signaling in mammals promotes the progression of ductal tumors (Yeh, Hou et al. 2006, Tang, Sun et al. 2013, Insua-Rodriguez, Pein et al. 2018). Here we identify Idgf3 as an additional component that feeds into JNK signaling. Ultimately in Ras^V12^-expressing SGs this leads to the formation of EnVs involving Spectrins. Under physiological conditions, members of the Spectrin family have a supporting role in maintaining cellular architecture through interaction with phospholipids and actively promoting polymerization of F-actin (Juliano, Kimelberg et al. 1971, Pinder, Bray et al. 1975, Hardy and Schrier 1978). Moreover, the secretory activity of ductal organs has been shown to be facilitated by Spectrins (Lattner, Leng et al. 2019).

During *Drosophila* development and under physiological conditions, the pathway that involves Spectrins, Rac1 and Pak1 has been shown to be required for the maintenance of cell polarity while when deregulated it leads to the formation of enlarged vesicles similar to the EnVs (Lee and Thomas 2011). Thus, our results provide a possible link between the observed induction of CLPs in a range of tumors and the effects of Spectrins and their deregulation in tumors (Ackermann and Brieger 2019, Yang, Yang et al. 2021). In addition to the genetic interaction we find, previous work suggests an additional mechanical link via a Spectrin binding protein (Human spectrin Src homology domain binding protein1; Hssh3bp1, (Ziemnicka-Kotula, Xu et al. 1998)) the loss of which has been associated with prostatic tumors (Macoska, Xu et al. 2001). Hhh3bp1 may influence tumor progression possibly through interaction with tyrosine kinases such as Abelson kinase (Macoska, Xu et al. 2001). Interestingly Hhh3bp1 is a marker and possible regulator of macropinocytosis (Dubielecka, Cui et al. 2010), a recycling pathway that is known to be hijacked by Ras-transformed tumor cells to acquire nutrients (Recouvreux and Commisso 2017) and also leads to the formation of large intracellular vesicles (Ritter, Bresgen et al. 2021). In favor of this hypothesis macropinocytosis is known to depend on Rac1/Pak1 signaling although the resulting vesicles are usually smaller (0.2-5 micrometers) than EnVs (Maxson, Sarantis et al. 2021). We find that - like macropinocytosis - EnV-formation depends on the activity of growth factors (Recouvreux and Commisso 2017), in this case Idgf3, much in line with its original description as an *in vitro* mediator of insulin signaling (Kawamura, Shibata et al. 1999). *In vivo*, under normal conditions Idgf3 is required for proper formation of chitin-containing structures, wound healing and cellular integrity (Pesch, Riedel et al. 2016). Thus, under these circumstances Idgf3 acts to preserve cellular integrity including the epithelial character of SG cells upstream of spectrins. Conversely, in a non-physiological setting such as upon overexpression of *Ras^V12^*, this mechanism is overwhelmed leading to the breakdown of homeostasis, loss of cell polarity and the gland lumen, loss of secretory activity and the formation of EnVs larger than macropinocytotic vesicles. Large vesicles accompany several scenarios of non-apoptotic programed cell death, which occurs a.o. in apoptosis-resistant tumors (Shubin, Demidyuk et al. 2016, Yan, Dawood et al. 2020). Such modes of cell death include methuosis, a deregulated form of macropinocytosis (Shubin, Demidyuk et al. 2016, Ritter, Bresgen et al. 2021). Of note, apoptotic cell death is inhibited in *Drosophila* polytenic SGs to account for the increased number of DNA breaks that occur during endoreplication, which in mitotic cells induce apoptosis in both a p53-dependent and independent manner (Mehrotra, Maqbool et al. 2008, Zhang, Mehrotra et al. 2014). In line, despite the activation of caspase activity and nuclear fragmentation, which are considered hallmarks of apoptosis, *Ras^V12^* SG cells don’t disintegrate to produce apoptotic bodies (Krautz, Khalili et al. 2020). This may also explain the difference to mitotically cycling tumor models, which also activate JNK – yet with apoptosis as an outcome (Uhlirova and Bohmann 2006, Araki, Kurihara et al. 2019, Parvy, Yu et al. 2019). Thus, SGs provide a suitable model for apoptosis-resistant tumors. In a mammalian setting, the phenotypes that are associated with non-apoptotic cell death such as disruption of cellular polarity and reorganization of the ECM provide potential targets for therapeutic treatments (Insua-Rodriguez, Pein et al. 2018). Our work adds CLPs and spectrins to this list. Depending on the tissue environment and similar to JNK signaling, CLP’s may have varying roles in a context-dependent manner. Overexpression of *Idgf3* alone is not sufficient for the loss of cell polarity, overgrowth, and fibrosis. Collectively, this suggests a tumor-specific phenotype for Idgf3 (Fig. 6B-J), in line with mammalian CLPs (reviewed in (Zhao, Su et al. 2020)). Due to their pleiotropic effects, further investigation of CLPs role will be required to dissect their molecular function in a given tissue and to ultimately design tumor-specific treatments (Kzhyshkowska, Larionova et al. 2019).

Taken together our findings provide new insight into the loss of tissue integrity in a neoplastic tumor model including the contribution of CLPs, Spectrins and alternative forms of cell death. This may provide further ways to test how developmentally and physiologically important conserved mechanisms that maintain cellular hemostasis - when deregulated - contribute to tumor progression.

## Supporting information

Complete supplementary material

## 6 Author Contributions

D.K., U.T and M.K. conceived the research and designed the experiments; D.K., M.K., S.H. and A.M. performed experiments and data analysation, D.K., U.T. and M.K. wrote the paper and participated in the revisions. All authors read and approved the final manuscript.

## 7 Funding

Swedish Cancer Foundation (CAN 2015-546)

Wenner-Gren Foundation (UPD2020-0094 and UPD2021-0095 to MK)

Swedish Research Council (VR 2016-04077 and VR 2021-04841)

## 8 Acknowledgments

We would like to thank Chris Molenaar, Roger Karlsson, Stina Höglund and the Imaging facility at Stockholm University for support with all aspects of microscopy. We would also like to thank Vasilios Tsarouhas for his critical feedback. This work was supported by grants from the Swedish Cancer Foundation (CAN 2015-546), the Wenner-Gren Foundation (UPD2020-0094 and UPD2021-0095 to MK) and the Swedish Research Council (VR 2016-04077 and VR 2021-04841).

## Reference

Ackermann, A. and A. Brieger (2019). “The Role of Nonerythroid Spectrin alphaII in Cancer.” J Oncol 2019: 7079604.

Araki, M., M. Kurihara, S. Kinoshita, R. Awane, T. Sato, Y. Ohkawa and Y. H. Inoue (2019). “Anti-tumour effects of antimicrobial peptides, components of the innate immune system, against haematopoietic tumours in Drosophila mxc mutants.” Dis Model Mech 12(6).

Archibald, A., C. Mihai, I. G. Macara and L. McCaffrey (2015). “Oncogenic suppression of apoptosis uncovers a Rac1/JNK proliferation pathway activated by loss of Par3.” Oncogene 34(24): 3199–3206.

Baek, S. H., Y. C. Kwon, H. Lee and K. M. Choe (2010). “Rho-family small GTPases are required for cell polarization and directional sensing in Drosophila wound healing.” Biochem Biophys Res Commun 394(3): 488–492.

Bartlett, B. J., P. Isakson, J. Lewerenz, H. Sanchez, R. W. Kotzebue, R. C. Cumming, G. L. Harris, I. P. Nezis, D. R. Schubert, A. Simonsen and K. D. Finley (2011). “p62, Ref(2)P and ubiquitinated proteins are conserved markers of neuronal aging, aggregate formation and progressive autophagic defects.” Autophagy 7(6): 572–583.

Bennett, V. and A. J. Baines (2001). “Spectrin and ankyrin-based pathways: metazoan inventions for integrating cells into tissues.” Physiol Rev 81(3): 1353–1392.

Brumby, A. M. and H. E. Richardson (2003). “scribble mutants cooperate with oncogenic Ras or Notch to cause neoplastic overgrowth in Drosophila.” EMBO J 22(21): 5769–5779.

Ciapponi, L., D. B. Jackson, M. Mlodzik and D. Bohmann (2001). “Drosophila Fos mediates ERK and JNK signals via distinct phosphorylation sites.” Genes Dev 15(12): 1540–1553.

Diwanji, N. and A. Bergmann (2017). “The beneficial role of extracellular reactive oxygen species in apoptosis-induced compensatory proliferation.” Fly (Austin) 11(1): 46–52.

Dubielecka, P. M., P. Cui, X. Xiong, S. Hossain, S. Heck, L. Angelov and L. Kotula (2010). “Differential regulation of macropinocytosis by Abi1/Hssh3bp1 isoforms.” PLoS One 5(5): e10430.

Fletcher, G. C., A. Elbediwy, I. Khanal, P. S. Ribeiro, N. Tapon and B. J. Thompson (2015). “The Spectrin cytoskeleton regulates the Hippo signalling pathway.” EMBO J 34(7): 940–954.

Fogarty, C. E. and A. Bergmann (2017). “Killers creating new life: caspases drive apoptosis-induced proliferation in tissue repair and disease.” Cell Death Differ 24(8): 1390–1400.

Hardy, B. and S. L. Schrier (1978). “The role of spectrin in erythrocyte ghost endocytosis.” Biochem Biophys Res Commun 81(4): 1153–1161.

Igaki, T., R. A. Pagliarini and T. Xu (2006). “Loss of cell polarity drives tumor growth and invasion through JNK activation in Drosophila.” Curr Biol 16(11): 1139–1146.

Insua-Rodriguez, J., M. Pein, T. Hongu, J. Meier, A. Descot, C. M. Lowy, E. De Braekeleer, H. P. Sinn, S. Spaich, M. Sutterlin, A. Schneeweiss and T. Oskarsson (2018). “Stress signaling in breast cancer cells induces matrix components that promote chemoresistant metastasis.” EMBO Mol Med 10(10).

Johansen, J. S., B. V. Jensen, A. Roslind, D. Nielsen and P. A. Price (2006). “Serum YKL-40, a new prognostic biomarker in cancer patients?” Cancer Epidemiol Biomarkers Prev 15(2): 194–202.

Juliano, R. L., H. K. Kimelberg and D. Papahadjopoulos (1971). “Synergistic effects of a membrane protein (spectrin) and Ca 2+ on the Na + permeability of phospholipid vesicles.” Biochim Biophys Acta 241(3): 894–905.

Karlsson, C., A. M. Korayem, C. Scherfer, O. Loseva, M. S. Dushay and U. Theopold (2004). “Proteomic analysis of the Drosophila larval hemolymph clot.” J Biol Chem 279(50): 52033–52041.

Kawamura, K., T. Shibata, O. Saget, D. Peel and P. J. Bryant (1999). “A new family of growth factors produced by the fat body and active on Drosophila imaginal disc cells.” Development 126(2): 211–219.

Khalili, D., C. Kalcher, S. Baumgartner and U. Theopold (2021). “Anti-Fibrotic Activity of an Antimicrobial Peptide in a Drosophila Model.” J Innate Immun 13(6): 376–390.

Khalili, D., C. Kalcher, S. Baumgartner and U. Theopold (2021). “Anti-Fibrotic Activity of an Antimicrobial Peptide in a Drosophila Model.” J Innate Immun: 1–15.

Kirkpatrick, R. B., R. E. Matico, D. E. McNulty, J. E. Strickler and M. Rosenberg (1995). “An abundantly secreted glycoprotein from Drosophila melanogaster is related to mammalian secretory proteins produced in rheumatoid tissues and by activated macrophages.” Gene 153(2): 147–154.

Krautz, R., D. Khalili and U. Theopold (2020). “Tissue-autonomous immune response regulates stress signalling during hypertrophy.” Elife 9.

Kucerova, L., V. Broz, B. Arefin, H. O. Maaroufi, J. Hurychova, H. Strnad, M. Zurovec and U. Theopold (2015). “The Drosophila Chitinase-Like Protein IDGF3 Is Involved in Protection against Nematodes and in Wound Healing.” J Innate Immun.

Kucerova, L., O. I. Kubrak, J. M. Bengtsson, H. Strnad, S. Nylin, U. Theopold and D. R. Nassel (2016). “Slowed aging during reproductive dormancy is reflected in genome-wide transcriptome changes in Drosophila melanogaster.” BMC Genomics 17(1): 50.

Kzhyshkowska, J., I. Larionova and T. Liu (2019). “YKL-39 as a Potential New Target for Anti-Angiogenic Therapy in Cancer.” Front Immunol 10: 2930.

Lattner, J., W. Leng, E. Knust, M. Brankatschk and D. Flores-Benitez (2019). “Crumbs organizes the transport machinery by regulating apical levels of PI(4,5)P2 in Drosophila.” Elife 8.

Lee, C. G., C. A. Da Silva, C. S. Dela Cruz, F. Ahangari, B. Ma, M. J. Kang, C. H. He, S. Takyar and J. A. Elias (2011). “Role of chitin and chitinase/chitinase-like proteins in inflammation, tissue remodeling, and injury.” Annual review of physiology 73: 479–501.

Lee, S. K. and G. H. Thomas (2011). “Rac1 modulation of the apical domain is negatively regulated by beta (Heavy)-spectrin.” Mech Dev 128(1-2): 116–128.

Macoska, J. A., J. Xu, D. Ziemnicka, T. S. Schwab, M. A. Rubin and L. Kotula (2001). “Loss of expression of human spectrin src homology domain binding protein 1 is associated with 10p loss in human prostatic adenocarcinoma.” Neoplasia 3(2): 99–104.

Maxson, M. E., H. Sarantis, A. Volchuk, J. H. Brumell and S. Grinstein (2021). “Rab5 regulates macropinocytosis by recruiting the inositol 5-phosphatases OCRL and Inpp5b that hydrolyse PtdIns(4,5)P2.” J Cell Sci 134(7).

Mehrotra, S., S. B. Maqbool, A. Kolpakas, K. Murnen and B. R. Calvi (2008). “Endocycling cells do not apoptose in response to DNA rereplication genotoxic stress.” Genes Dev 22(22): 3158–3171.

Morera, E., S. S. Steinhauser, Z. Budkova, S. Ingthorsson, J. Kricker, A. Krueger, G. A. Traustadottir and T. Gudjonsson (2019). “YKL-40/CHI3L1 facilitates migration and invasion in HER2 overexpressing breast epithelial progenitor cells and generates a niche for capillary-like network formation.” In Vitro Cell Dev Biol Anim 55(10): 838–853.

Pagliarini, R. A. and T. Xu (2003). “A genetic screen in Drosophila for metastatic behavior.” Science 302(5648): 1227–1231.

Palmerini, V., S. Monzani, Q. Laurichesse, R. Loudhaief, S. Mari, V. Cecatiello, V. Olieric, S. Pasqualato, J. Colombani, D. S. Andersen and M. Mapelli (2021). “Drosophila TNFRs Grindelwald and Wengen bind Eiger with different affinities and promote distinct cellular functions.” Nat Commun 12(1): 2070.

Park, K. R., H. M. Yun, K. Yoo, Y. W. Ham, S. B. Han and J. T. Hong (2020). “Chitinase 3 like 1 suppresses the stability and activity of p53 to promote lung tumorigenesis.” Cell Commun Signal 18(1): 5.

Parvy, J. P., Y. Yu, A. Dostalova, S. Kondo, A. Kurjan, P. Bulet, B. Lemaitre, M. Vidal and J. B. Cordero (2019). “The antimicrobial peptide defensin cooperates with tumour necrosis factor to drive tumour cell death in Drosophila.” Elife 8.

Perez, E., J. L. Lindblad and A. Bergmann (2017). “Tumor-promoting function of apoptotic caspases by an amplification loop involving ROS, macrophages and JNK in Drosophila.” Elife 6.

Pesch, Y. Y., D. Riedel, K. R. Patil, G. Loch and M. Behr (2016). “Chitinases and Imaginal disc growth factors organize the extracellular matrix formation at barrier tissues in insects.” Sci Rep 6: 18340.

Pinder, J. C., D. Bray and W. B. Gratzer (1975). “Actin polymerisation induced by spectrin.” Nature 258(5537): 765–766.

Recouvreux, M. V. and C. Commisso (2017). “Macropinocytosis: A Metabolic Adaptation to Nutrient Stress in Cancer.” Front Endocrinol (Lausanne) 8: 261.

Ritter, M., N. Bresgen and H. H. Kerschbaum (2021). “From Pinocytosis to Methuosis-Fluid Consumption as a Risk Factor for Cell Death.” Front Cell Dev Biol 9: 651982.

Roslind, A. and J. S. Johansen (2009). “YKL-40: a novel marker shared by chronic inflammation and oncogenic transformation.” Methods Mol Biol 511: 159–184.

Shao, R., K. Hamel, L. Petersen, Q. J. Cao, R. B. Arenas, C. Bigelow, B. Bentley and W. Yan (2009). “YKL-40, a secreted glycoprotein, promotes tumor angiogenesis.” Oncogene 28(50): 4456–4468.

Shubin, A. V., I. V. Demidyuk, A. A. Komissarov, L. M. Rafieva and S. V. Kostrov (2016). “Cytoplasmic vacuolization in cell death and survival.” Oncotarget 7(34): 55863–55889.

Tang, H., Y. Sun, Z. Shi, H. Huang, Z. Fang, J. Chen, Q. Xiu and B. Li (2013). “YKL-40 induces IL-8 expression from bronchial epithelium via MAPK (JNK and ERK) and NF-kappaB pathways, causing bronchial smooth muscle proliferation and migration.” J Immunol 190(1): 438–446.

Uhlen, M., C. Zhang, S. Lee, E. Sjostedt, L. Fagerberg, G. Bidkhori, R. Benfeitas, M. Arif, Z. Liu, F. Edfors, K. Sanli, K. von Feilitzen, P. Oksvold, E. Lundberg, S. Hober, P. Nilsson, J. Mattsson, J. M. Schwenk, H. Brunnstrom, B. Glimelius, T. Sjoblom, P. H. Edqvist, D. Djureinovic, P. Micke, C. Lindskog, A. Mardinoglu and F. Ponten (2017). “A pathology atlas of the human cancer transcriptome.” Science 357(6352).

Uhlirova, M. and D. Bohmann (2006). “JNK- and Fos-regulated Mmp1 expression cooperates with Ras to induce invasive tumors in Drosophila.” Embo J 25(22): 5294–5304.

Wertheimer, E., A. Gutierrez-Uzquiza, C. Rosemblit, C. Lopez-Haber, M. S. Sosa and M. G. Kazanietz (2012). “Rac signaling in breast cancer: a tale of GEFs and GAPs.” Cell Signal 24(2): 353–362.

Williams, S. T., A. N. Smith, C. D. Cianci, J. S. Morrow and T. L. Brown (2003). “Identification of the primary caspase 3 cleavage site in alpha II-spectrin during apoptosis.” Apoptosis 8(4): 353–361.

Yadav, S. and I. Eleftherianos (2018). “The Imaginal Disc Growth Factors 2 and 3 participate in the Drosophila response to nematode infection.” Parasite Immunol 40(10): e12581.

Yan, G., M. Dawood, M. Bockers, S. M. Klauck, C. Fottner, M. M. Weber and T. Efferth (2020). “Multiple modes of cell death in neuroendocrine tumors induced by artesunate.” Phytomedicine 79: 153332.

Yang, P., Y. Yang, P. Sun, Y. Tian, F. Gao, C. Wang, T. Zong, M. Li, Y. Zhang, T. Yu and Z. Jiang (2021). “betaII spectrin (SPTBN1): biological function and clinical potential in cancer and other diseases.” Int J Biol Sci 17(1): 32–49.

Yeh, Y. T., M. F. Hou, Y. F. Chung, Y. J. Chen, S. F. Yang, D. C. Chen, J. H. Su and S. S. Yuan (2006). “Decreased expression of phosphorylated JNK in breast infiltrating ductal carcinoma is associated with a better overall survival.” Int J Cancer 118(11): 2678–2684.

Zeke, A., M. Misheva, A. Remenyi and M. A. Bogoyevitch (2016). “JNK Signaling: Regulation and Functions Based on Complex Protein-Protein Partnerships.” Microbiol Mol Biol Rev 80(3): 793–835.

Zhang, B., S. Mehrotra, W. L. Ng and B. R. Calvi (2014). “Low levels of p53 protein and chromatin silencing of p53 target genes repress apoptosis in Drosophila endocycling cells.” PLoS Genet 10(9): e1004581.

Zhao, T., Z. Su, Y. Li, X. Zhang and Q. You (2020). “Chitinase-3 like-protein-1 function and its role in diseases.” Signal Transduct Target Ther 5(1): 201.

Zhu, M., T. Xin, S. Weng, Y. Gao, Y. Zhang, Q. Li and M. Li (2010). “Activation of JNK signaling links lgl mutations to disruption of the cell polarity and epithelial organization in Drosophila imaginal discs.” Cell Res 20(2): 242–245.

Ziemnicka-Kotula, D., J. Xu, H. Gu, A. Potempska, K. S. Kim, E. C. Jenkins, E. Trenkner and L. Kotula (1998). “Identification of a candidate human spectrin Src homology 3 domain-binding protein suggests a general mechanism of association of tyrosine kinases with the spectrin-based membrane skeleton.” J Biol Chem 273(22): 13681–13692.

