## Supplementary material for "Chitinase-like proteins promoting tumorigenesis through disruption of cell polarity via enlarged endosomal vesicles": Complete supplementary material

### Chitinase-like proteins promote tumorigenesis through disruption of cell polarity and formation of enlarged vesicles

#### 1 Supplementary figures

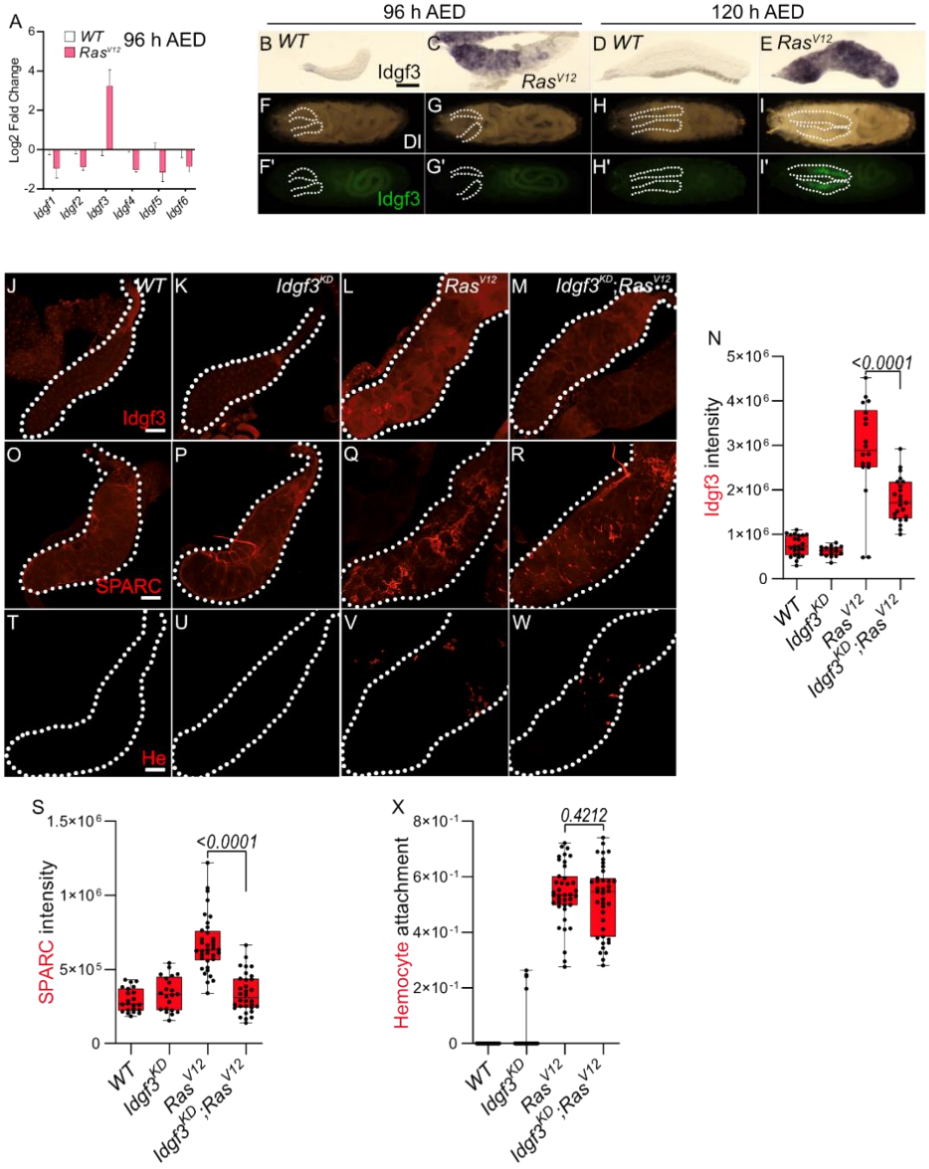

**Figure S1**
**Idgf3 characterization and tumor effects**
**(A)** qPCR data showing induction of *Idgf3* in 96 h AED $Ras^{V12}$ glands. **(B-E)** ISH showing Idgf3 distribution throughout the SG at 96 h and 120 h AED. **(F-I')** Whole larvae images showing Idgf3::GFP localization. **(J-M)** Antibody staining against Idgf3 showing efficiency of $Idgf3^{KD}$. **(N)** Intensity quantification showing $Idgf3^{KD}$ reduction efficiency of Idgf3. **(O-R)** SPARC staining displaying reduced fibrosis in $Idgf3^{KD};Ras^{V12}$ SG. **(S)** Quantification showing reduced SPARC intensity in $Idgf3^{KD};Ras^{V12}$ SG. **(T-W)** Hemocytes staining. **(X)** Quantification showing no effect on attached hemocytes in $Idgf3^{KD};Ras^{V12}$ SG. Data in **(A)** represent 3 independent replicas summarized as mean ± SD Scale bars in **(B-E)** represent 0.3 mm and **(J-W)** 100 µm. Boxplot in **(N, S and X)** represent at least 20 SG pairs. Whisker length min to max, bar represent median. P-value quantified with Student's t-test.

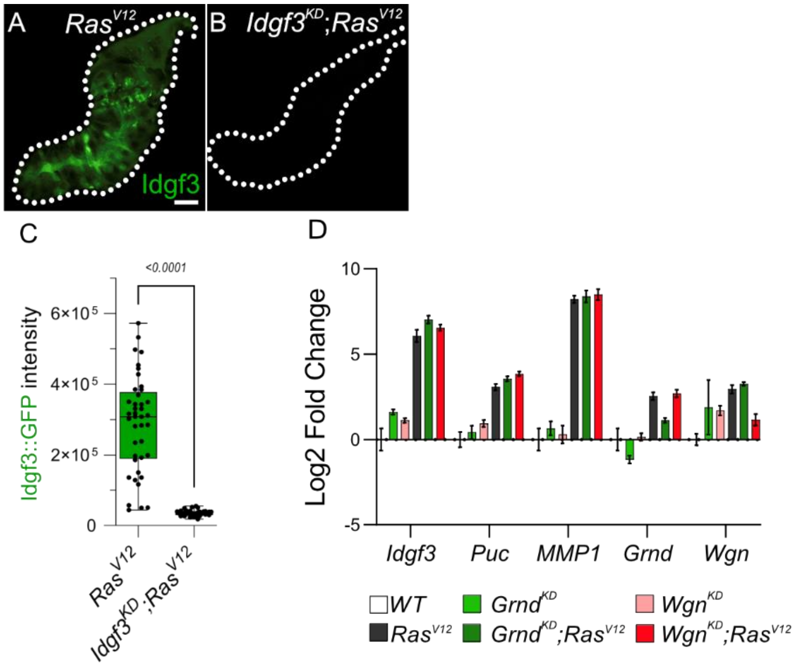


**Figure S2**
**Non-canonical JNK regulation and activation in $Ras^{V12}$ glands**
**(A-B)** $Idgf3^{KD};Ras^{V12}$ showing reduced Idgf3::GFP intensity quantified in **(C)**. **(D)** qPCR showing $Grnd^{KD}$ and $Wgn^{KD}$ are downregulating respective mRNA efficiently in $Ras^{V12}$ background. Scale bars in **(A-B)** represent 100 µm. Boxplot in **(C)** represent at least 20 SG pairs. Whisker length min to max, bar represent median. Data in **(D)** represent 4 independent replicas summarized as mean ± SD. P-value quantified with Student's t-test.

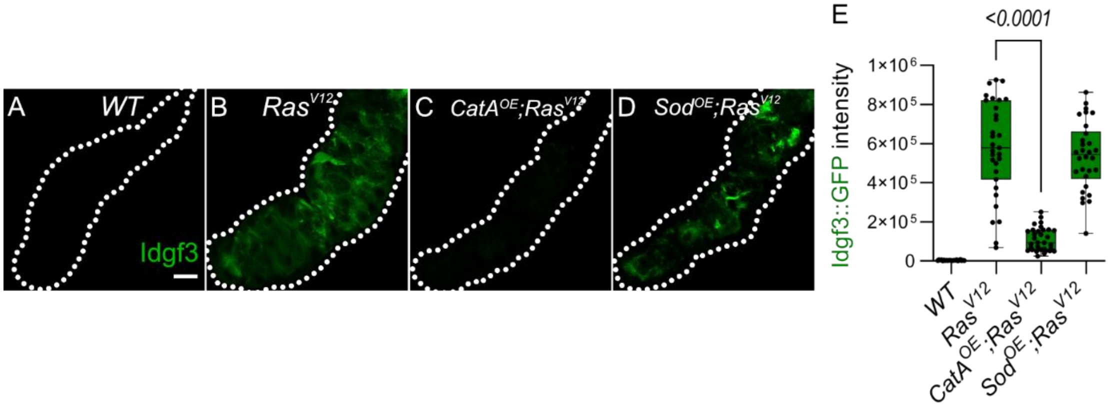


**Figure S3**

**Idgf3 and JNK regulation by ROS**

**(A-D)** Effect on Idgf3::GFP by reductases. **(E)** Quantification showing reduced Idgf3::GFP intensity in $CatA^{OE};Ras^{V12}$. Scale bars in **(A-D)** represent 100 µm. Boxplot in **(E)** represent at least 20 SG pairs. Whisker length min to max, bar represent median. P-value quantified with Student's t-test.

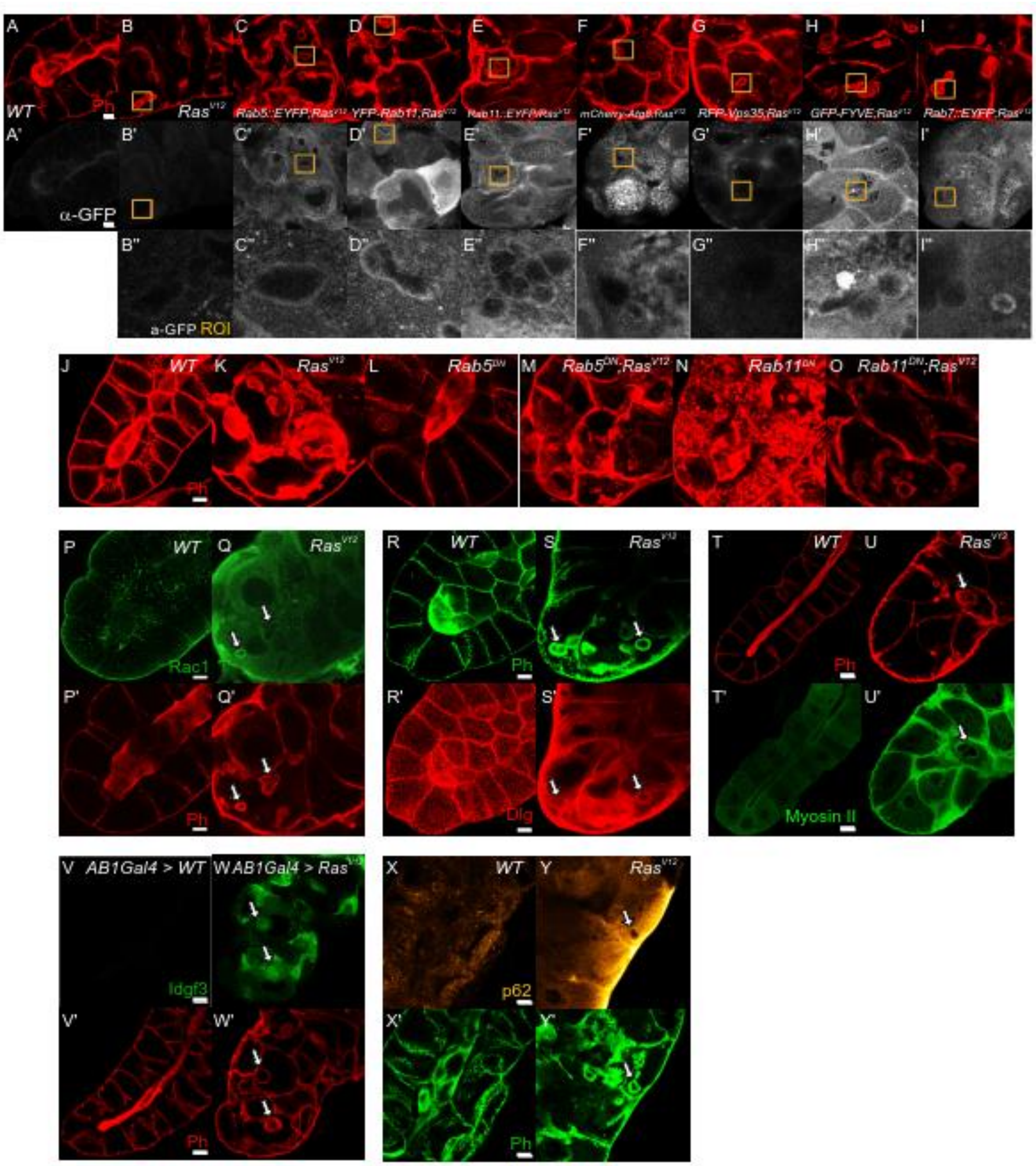


**Figure S4**

**Enlarged vesicles (EnVs) characterization**

**(A-I'')** Vesicle markers showing EnVs stained for RAb5, Rab11 and FYVE. **(J-O)** Phalloidin staining showing no effect on the formation of EnVs in *Rab5*$^{DN}$;*Ras*$^{V12}$ and *Rab11*$^{DN}$;*Ras*$^{V12}$ glands. **(P-Q')** Rac1 staining showing EnVs co-localizing with Phalloidin. **(R-S')** Phalloidin and Dlg staining showing to co-localize at EnVs. **(T-U')** Phalloidin and GFP tagged Myosin II showing to co-localize at EnVs. **(V-W')** Idgf3 induction and accumulation are independent of the SG driver. **(X-Y')** p62 staining showing no accumulation in EnVs. Scale bars in **(A-Y')** represent 20 µm.

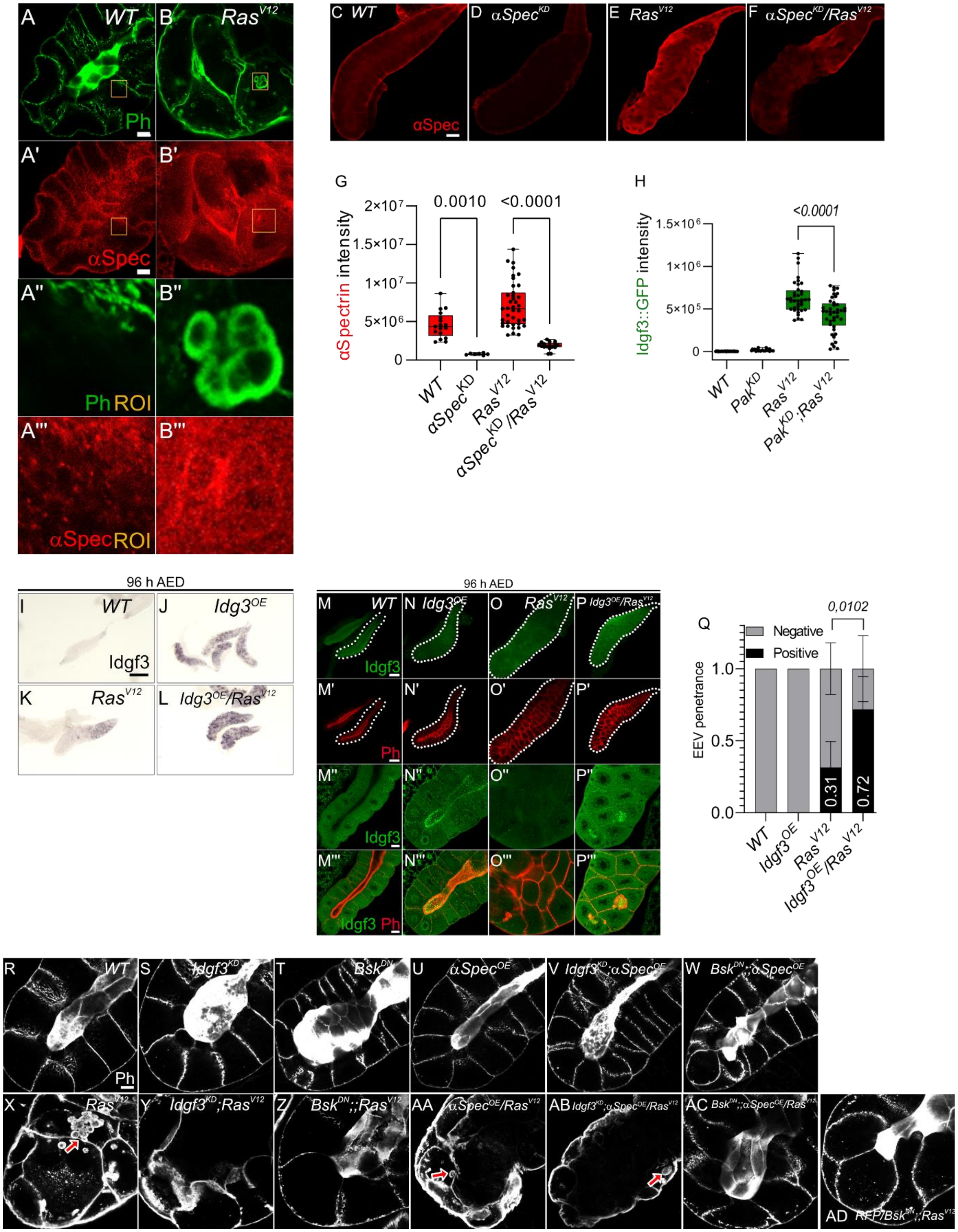


**Figure S5**

**(A-B'')** Phalloidin and αSpectrin staining shows colocalization at EnVs. **(C-F)** αSpectrin staining showing reduction in $\alpha Spectrin^{KD}$ quantified in **(G)**. **(H)** Quantification showing increased level of Idgf3::GFP, no formation of EnVs and Idgf3 in $Pak^{KD};Ras^{V12}$ glands. **(I-L)** ISH showing Idgf3 distribution in $Idgf3^{OE}/Ras^{V12}$ glands. **(M-P''')** Idgf3 staining EnVs in $Idgf3^{OE};Ras^{V12}$ glands. **(Q)**

EnV penetrance quantification showing increased amount of glands with EnVs in Idgf3$^{OE}$;Ras$^{V12}$. **(R-AD)** Epistatic analysis between Idgf3 and αSpectrin. Phalloidin staining showing EnVs presence in *Idgf3$^{KD}$;αSpec$^{OE}$/ Ras$^{V12}$*. Scale bars in **(A-B‘, M‘‘-P‘‘‘, R-AD)** represent 20 µm, (C-F, M-P‘) represents 100 µm, **(I-L)** represents 0.3 mm. Boxplot in **(G-H)** represent at least 20 SG pairs. Bar plot in **(Q)** represent 3 independent replicas with at least 10 SGs pairs summarized as relative value ± SD. Whisker length min to max, bar represent median. P-value quantified with Student’s t-test.

#### 2 Supplementary tables

##### Supplementary table 1.

| REAGENT or RESOURCE | SOURCE | IDENTIFIER |
|---|---|---|
| **Antibodies** | | |
| anti-IDGF3 (rabbit) | Kucerova et al., 2016 | Idgf3 |
| anti-SPARC | Khalili et al., 2021 | Sparc |
| anti-Hemese (mouse monoclonal) | István Andó | H2 |
| pJNK | Cell Signaling Technology | 9255 |
| Rac1 fitc conjugated mouse | BD Bioscience Pharmingen | 610652 |
| αSpectrin | DSHB | 3A9 |
| anti-GFP (mouse monoclonal) | ThemoFisher Scientific | A11120 |
| anti-mouse-IgG Alexa546 (goat polyclonal) | ThemoFisher Scientific | A11030 |
| anti-rabbit-IgG Alexa568 (goat polyclonal) | ThemoFisher Scientific | A21069 |
| anti-mouse-IgG, Alexa Fluor 488 | ThemoFisher Scientific | A11001 |
| anti-rabbit-IgG, Alexa Fluor 488 | ThemoFisher Scientific | A11008 |
| **Chemicals, peptides, and recombinant proteins** | | |
| DAPI | Sigma-Aldrich | D9542 |
| Alexa Fluor 488 Phalloidin | ThermoFisher Scientific | A12379 |
| Alexa Fluor 546 Phalloidin | ThermoFisher Scientific | A22283 |
| Vancomycin | Sigma-Aldrich | V2002 |
| Neomycin | Sigma-Aldrich | N1876 |
| Metronidazole | Sigma-Aldrich | M3761 |
| Carbenicillin | Sigma-Aldrich | C1389 |
| **Critical commercial assays** | | |
| RNAquesous Micro Kit | ThemoFisher Scientific | AM1931 |
| **Experimental models: organisms/strains** | | |
| *D. melanogaster w$^{1118}$* | | |
| *D. melanogaster Bx$^{ms1096}$* | BDSC | 8860 |
| *D. melanogaster UAS-Ras$^{V12}$* | BDSC | 4847 |
| *D. melanogaster Idgf3::GFP* | Kucerova et al., 2016 | |
| *D. melanogaster UAS-Idgf3$^{KD}$* | BDSC | |
| *D. melanogaster UAS-Grnd$^{KD}$* | VDRC | 104538 |
| *D. melanogaster UAS-Wgn$^{KD}$* | BDSC | 55275 |
| *D. melanogaster UAS-Bsk$^{KD}$* | BDSC | 36643 |

| | | |
|---|---|---|
| *D. melanogaster UAS-Bsk*$^{KD}$ | VDRC | 104569 |
| *D. melanogaster UAS-Drs* | B. Lemaitre | |
| *D. melanogaster UAS-IRC* | WJL | |
| *D. melanogaster UAS- CatA* | BDSC | 24621 |
| *D. melanogaster UAS-SodA* | BDSC | 33605 |
| *D. melanogaster UAS-MFG-E8* | Dr Nakanishi | |
| *D. melanogaster UAS-MFG-E8ΔC* | Dr Nakanishi | |
| *D. melanogaster Rab5::EYFP* | BDSC | 62543 |
| *D. melanogaster UAS-YFP.Rab11* | BDSC | 50782 |
| *D. melanogaster Rab7::EYFP* | BDSC | 62545 |
| *D. melanogaster Rab11::EYFP* | BDSC | 62549 |
| *D. melanogaster UAS-mCherry-Atg8* | BDSC | 37750 |
| *D. melanogaster RFP-Vps35* | BDSC | 66527 |
| *D. melanogaster GFP-FYVE* | BDSC | 42712 |
| *D. melanogaster UAS-YFP.Rab5*$^{DN}$ | BDSC | 9771 |
| *D. melanogaster UAS-YFP.Rab11*$^{DN}$ | BDSC | 9792 |
| *D. melanogaster UAS-Rac1* | BDSC | 6293 |
| *D. melanogaster UAS-Pak*$^{KD}$ | BDSC | 41714 |
| *D. melanogaster UAS-βSpect*$^{KD}$ | BDSC | |
| *D. melanogaster UAS-αSpec*$^{KD}$ | BDSC | 31209 |
| *D. melanogaster UAS-Idgf3* | Kucerova et al., 2016<br>BDSC | 52658 |
| *D. melanogaster mCD8::RFP* | Krautz et al., 2020<br>BDSC | 27400 |
| *D. melanogaster αSpec.UPS* | BDSC | 32005 |
| *D. melanogaster TRE.GFP* | D. Bohmann | |
| *D. melanogaster hs-Flp*$^{122}$*; UAS-mCherry/CyO, Act-GFP*$^{JMR1}$*; UAS-Flp, Act5C > CD2 > Gal4/TM6b* | Duan et al., 2020 | |
| **Oligonucleotides** | | |
| *Drosophila Idgf1* forward: 5'- CTCAGTGTGACAAGTTTGACCC-3' | | |
| *Drosophila Idgf1* reverse: 5'- CCTTGGCGCAAATACGAGTTG-3' | | |
| *Drosophila Idgf2* forward: 5'-CAAGAAAGGTTCACGGTGATCT-3' | | |
| *Drosophila Idgf2* reverse: 5'-AACTGCTCCTTATGAAGGGCT-3' | | |
| *Drosophila Idgf3* forward: 5'-GTGCCGCTCTGAAACAAAAT-3' | | |
| *Drosophila Idgf3* reverse: 5'-ACGGGGCATCATAGTACCAG-3' | | |
| *Drosophila Idgf4* forward: 5'-TTACTATGACGGCAACAGTTTTGT-3' | | |
| *Drosophila Idgf4* reverse: 5'-GGCATATCCGTAGACCAGATG-3' | | |
| *Drosophila Idgf5* forward: 5'-CAGAGGTTGGAAAACTGGTGT-3' | | |
| *Drosophila Idgf5* reverse: 5'-GTTCAGCCAGCGACATTTGG-3' | | |
| *Drosophila Idgf6* forward: 5'-ACCGCTCTACTCCGTGATGT-3' | | |
| *Drosophila Idgf6* reverse: 5'-GCGGGAATGTCGTAGAACAG-3' | | |
| *Drosophila Puc* forward: 5'-CGTCATCATCAACGGCAAT-3' | | |
| *Drosophila Puc* reverse: 5'-AGGCGGGGTGTGTTTCTAT-3' | | |
| *Drosophila MMP1* forward: 5'-GTTTCCACCACCACACAGG-3' | | |
| *Drosophila MMP1* reverse: 5'-GCAGAGGCGGGTAGATAGC-3' | | |
| *Drosophila Grnd* forward: 5'-CACAACTACGATGCGTTTCTGT-3' | | PP32778 |
| *Drosophila Grnd* reverse: 5'-CATCTCGGCTTTTAACGGCTC-3' | | PP32778 |
| *Drosophila Wng* forward: 5'-ACCATCTGCGGTTCCATATACG-3' | | PP26269 |
| *Drosophila Wgn* reverse: 5'-GTGCTCATACTCGGAGGACTT-3' | | PP26269 |

| | | |
|---|---|---|
| *Drosophila Hid* forward: 5'-TCTACGAGTGGGTCAGGATGT-3' | | |
| *Drosophila Hid* reverse: 5'-GCGGATACTGGAAGATTTGC-3' | | |
| *Drosophila Rac1* forward: 5'-GGAAATCGAACCATGCAGGC-3' | | PD70033 |
| *Drosophila Rac1* reverse: 5'-GTCGAACACGGTGGGTATGT-3' | | PD70033 |
| *Drosophila αSpec* forward: 5'-CGACCGCCCCTATGTAACTA-3' | | PD70352 |
| *Drosophila αSpec* reverse: 5'-CACGCAGTAGTCAGCCATGT-3' | | PD70352 |
| *Drosophila Karst* forward: 5'-GGACAACCTTAACCATGCCTT-3' | | PP36727 |
| *Drosophila Karst* reverse: 5'-AGTAGGAGGCCACATAGGTCA-3' | | PP36727 |
| *Human CH3L1* forward: 5'-TCAAGAACAGGAACCCCAAC-3' | | |
| *Human CH3L1* reverse: 5'-AAATTCGGCCTTCATTTCCT -3' | | |
| *Human CH3L1* forward: 5'-CAGGGAGGCAAATGATTGAT -3' | | |
| *Human CH3L1* reverse: 5'-CCACCTTCTCTGATGGCATT -3' | | |
| *Drosophila Rpl32* forward: 5'-CGGATCGATATGCTA-3' | | |
| *Drosophila Rpl32* reverse: 5'-CGACGCACTCTGTTG-3' | | |
| **Recombinant DNA** | | |
| Idgf3 cDNA | Drosophila Genomics Resource Center (DGRC) | GH07453 |
| **Software and algorithms** | | |
| FIJI (ImageJ) | https://imagej.net/Fiji/Downloads | Version 1.53j |
| GraphPad Prism | https://www.graphpad.com/ | Version 9.1.2 |
| Affinity Designer | https://affinity.serif.com/en-gb/ | Version 1.9.2.1035 |
| Zen blue | https://www.zeiss.com/ | Version 2.3 |
| AxioVision LE | https://www.zeiss.com/ | Version 4.8.2.0 |

#### 2.1 Cross list:

**Fig. 1**

A:
♀ $w^{1118}$,*Beadex-Gal4 (Bx)* > ♂$w^{1118}$
♀*Bx* > ♂ $w^{1118}$;;*UAS-Ras*$^{V12}$ (*Ras*$^{V12}$)

B-C:
♀ *Bx; Idgf3::GFP* > ♂$w^{1118}$
♀ *Bx; Idgf3::GFP* > ♂ *Ras*$^{V12}$

D-Q:
♀*Bx* > ♂$w^{1118}$
♀*Bx* > ♂ *Ras*$^{V12}$

♀*Bx* > ♂ *w*$^{1118}$*; Idgf3*$^{KD}$
♀*Bx* > ♂ *w*$^{1118}$*; Idgf3*$^{KD}$*; Ras*$^{V12}$

**Fig. S1**
A:
♀*Bx* > ♂*w*$^{1118}$
♀*Bx* > ♂ *Ras*$^{V12}$

B-E:
♀*Bx* > ♂*w*$^{1118}$
♀*Bx* > ♂ *Ras*$^{V12}$

F-I':
♀ *Bx; Idgf3::GFP* > ♂*w*$^{1118}$
♀ *Bx; Idgf3::GFP* > ♂ *Ras*$^{V12}$

J-X:
♀*Bx* > ♂*w*$^{1118}$
♀*Bx* > ♂ *w*$^{1118}$*; Idgf3*$^{KD}$
♀*Bx* > ♂ *Ras*$^{V12}$
♀*Bx* > ♂ *w*$^{1118}$*; Idgf3*$^{KD}$*; Ras*$^{V12}$

**Fig. 2**
A-E:
♀ *Bx; Idgf3::GFP* > ♂ *Ras*$^{V12}$
♀ *Bx; Idgf3::GFP* > ♂ *w*$^{1118}$*; Grnd*$^{KD}$*; Ras*$^{V12}$
♀ *Bx; Idgf3::GFP* > ♂ *w*$^{1118}$*; Wgn*$^{KD}$*; Ras*$^{V12}$
♀ *Bx; Idgf3::GFP* > ♂ *w*$^{1118}$*; JNK*$^{KD\ [36643/Bl]}$*; Ras*$^{V12}$

**Fig. S2**
A-C:
♀ *Bx; Idgf3::GFP* > ♂ *Ras*$^{V12}$
♀ *Bx; Idgf3::GFP* > ♂ *w*$^{1118}$*; Idgf3*$^{KD}$*; Ras*$^{V12}$

D:
♀*Bx* > ♂*w*$^{1118}$
♀ *Bx* > ♂ *Ras*$^{V12}$
♀ *Bx* > ♂ *w*$^{1118}$*; Grnd*$^{KD}$
♀ *Bx* > ♂ *w*$^{1118}$*; Wgn*$^{KD}$
♀ *Bx* > ♂ *w*$^{1118}$*; Grnd*$^{KD}$*; Ras*$^{V12}$
♀ *Bx* > ♂ *w*$^{1118}$*; Wgn*$^{KD}$*; Ras*$^{V12}$

**Fig. 3**
A-I:
♀ *Bx* > ♂ *w*$^{1118}$*; Idgf3::GFP*
♀ *Bx;; IRC-OE* > ♂ *w*$^{1118}$*; Idgf3::GFP*
♀ *Bx* > ♂ *w*$^{1118}$*; Idgf3::GFP; Ras*$^{V12}$
♀ *Bx;; IRC-OE* > ♂ *w*$^{1118}$*; Idgf3::GFP; Ras*$^{V12}$

J-O:
♀ *Bx* > ♂ $w^{1118}$
♀ *Bx;; IRC-OE* > ♂ $w^{1118}$
♀ *Bx* > ♂ $Ras^{V12}$
♀ *Bx;; IRC-OE* > ♂ $Ras^{V12}$

P-T:
♀ *Bx* > ♂ $w^{1118}$; TRE.GFP/ CyO.GFP
♀ *Bx;; IRC-OE* > ♂ $w^{1118}$; TRE.GFP/ CyO.GFP
♀ *Bx* > ♂ $w^{1118}$; TRE.GFP/ CyO.GFP; $Ras^{V12}$
♀ *Bx;; IRC-OE* > ♂ $w^{1118}$; TRE.GFP/ CyO.GFP; $Ras^{V12}$

**Fig. S3**
A-E:
♀ *Bx; Idgf3::GFP* > ♂ $w^{1118}$
♀ *Bx; Idgf3::GFP* > ♂ $w^{1118}$; $Ras^{V12}$
♀ *Bx; Idgf3::GFP* > ♂ $w^{1118}$; *CatA-OE*; $Ras^{V12}$
♀ *Bx; Idgf3::GFP* > ♂ $w^{1118}$; *SodA-OE*; $Ras^{V12}$

**Fig. 4**
B-C':
♀*Bx; Idgf3::GFP*> ♂ $w^{1118}$
♀*Bx; Idgf3::GFP*> ♂ $Ras^{V12}$

D:
♀*Bx* > ♂ $w^{1118}$
♀*Bx* > ♂ $Ras^{V12}$

E-J':
♀ *Bx* > ♂ $w^{1118}$
♀ *Bx* > ♂ $w^{1118}$; *MFG-E8ΔC::GFP-OE*
♀ *Bx* > ♂ $w^{1118}$; *MFG-E8::GFP-OE*
♀ *Bx* > ♂ $Ras^{V12}$
♀ *Bx* > ♂ $w^{1118}$; *MFG-E8ΔC::GFP-OE*; $Ras^{V12}$
♀ *Bx* > ♂ $w^{1118}$; *MFG-E8::GFP-OE*; $Ras^{V12}$

**Fig. S4**
A-I'':
♀ *Bx* > ♂ $w^{1118}$
♀ *Bx* > ♂ $Ras^{V12}$
♀ *Bx* > ♂ $w^{1118}$; *Rab5::EYFP*; $Ras^{V12}$
♀ *Bx* > ♂ $w^{1118}$; *YFP-Rab11*; $Ras^{V12}$
♀ *Bx;; Rab11::EYFP* > ♂ $Ras^{V12}$
♀ *Bx* > ♂ $w^{1118}$; *GFP-FYVE*; $Ras^{V12}$
♀ *Bx;; Rab7::EYFP* > ♂ $w^{1118}$;; $Ras^{V12}$

J-O:
♀ *Bx* > ♂ $w^{1118}$
♀ *Bx* > ♂ $Ras^{V12}$
♀ *Bx* > ♂ $w^{1118}$*; Rab5*$^{DN}$
♀ *Bx* > ♂ $w^{1118}$*; Rab5*$^{DN}$*; Ras*$^{V12}$
♀ *Bx* > ♂ $w^{1118}$*; Rab11*$^{DN}$
♀ *Bx* > ♂ $w^{1118}$*; Rab11*$^{DN}$*; Ras*$^{V12}$

P-Q':
♀ *Bx* > ♂ $w^{1118}$
♀ *Bx* > ♂ $Ras^{V12}$

R-S':
♀ *Bx* > ♂ $w^{1118}$
♀ *Bx* > ♂ $Ras^{V12}$

T-U':
♀ *Bx;; sqh-GFP* > ♂ $w^{1118}$
♀ *Bx;; sqh-GFP* > ♂ $Ras^{V12}$

V-W':
♀ $w^{1118}$*; AB1-Gal4>* ♂$w^{1118}$*; Idgf3::GFP*
♀ $w^{1118}$*; AB1-Gal4>* ♂$w^{1118}$*; Idgf3::GFP; Ras*$^{V12}$

X-Y':
♀*Bx* > ♂$w^{1118}$
♀*Bx* > ♂ $Ras^{V12}$

**<u>Fig. 5</u>**
A-E:
♀*Bx* > ♂$w^{1118}$
♀*Bx* > ♂ $w^{1118}$*; Idgf3*$^{KD}$
♀*Bx* > ♂ $Ras^{V12}$
♀*Bx* > ♂ $w^{1118}$*; Idgf3*$^{KD}$*; Ras*$^{V12}$

F-J:
♀ *Bx* > ♂ $w^{1118}$*;; Idgf3::GFP*
♀ *Bx;; αSpec*$^{KD}$ > ♂ $w^{1118}$*; Idgf3::GFP*
♀ *Bx* > ♂ $w^{1118}$*; Idgf3::GFP; Ras*$^{V12}$
♀ *Bx;; αSpec*$^{KD}$ > ♂ $w^{1118}$*; Idgf3::GFP; Ras*$^{V12}$

K-T:
♀ *Bx* > ♂ $w^{1118}$
♀ *Bx* > ♂ $w^{1118}$*, JNK*$^{DN}$
♀ *Bx;; Idgf3-OE* > ♂ $w^{1118}$
♀ *Bx;; Idgf3-OE* > ♂ $w^{1118}$*, JNK*$^{DN}$
♀ *Bx* > ♂ $Ras^{V12}$
♀ *Bx* > ♂ $w^{1118}$*, JNK*$^{DN}$*;; Ras*$^{V12}$

♀ *Bx;; Idgf3-OE* > ♂ *Ras*$^{V12}$
♀ *Bx;; Idgf3-OE* > ♂ *w*$^{1118}$*, JNK*$^{DN}$*;; Ras*$^{V12}$
♀ *Bx, mCD8::RFP-OE* > ♂ *w*$^{1118}$*, JNK*$^{DN}$*;; Ras*$^{V12}$

U-V:
♀ *Bx* > ♂ *w*$^{1118}$*; TRE.GFP/CyO.GFP*
♀ *Bx;; αSpec*$^{KD}$ > ♂ *w*$^{1118}$*; TRE.GFP/CyO.GFP*
♀ *Bx* > ♂ *w*$^{1118}$*; TRE.GFP/CyO.GFP; Ras*$^{V12}$
♀ *Bx;; αSpec*$^{KD}$ > ♂ *w*$^{1118}$*; TRE.GFP/CyO.GFP; Ras*$^{V12}$

**Fig. S5**
A-B''':
♀ *Bx* > ♂ *w*$^{1118}$
♀ *Bx* > ♂ *Ras*$^{V12}$

C-G:
♀ *Bx* > ♂ *w*$^{1118}$*;; Idgf3::GFP*
♀ *Bx;; αSpec*$^{KD}$ > ♂ *w*$^{1118}$*; Idgf3::GFP*
♀ *Bx* > ♂ *w*$^{1118}$*; Idgf3::GFP; Ras*$^{V12}$
♀ *Bx;; αSpec*$^{KD}$ > ♂ *w*$^{1118}$*; Idgf3::GFP; Ras*$^{V12}$

H:
♀ *Bx; Idgf3::GFP* > ♂ *w*$^{1118}$
♀ *Bx; Idgf3::GFP* > ♂ *w*$^{1118}$*; Pak*$^{KD}$
♀ *Bx; Idgf3::GFP* > ♂ *Ras*$^{V12}$
♀ *Bx; Idgf3::GFP* > ♂ *w*$^{1118}$*; Pak*$^{KD}$*; Ras*$^{V12}$

I-Q:
♀Bx > ♂w$^{1118}$
♀*Bx;; Idgf3-OE* > ♂ *w*$^{1118}$
♀*Bx* > ♂ *Ras*$^{V12}$
♀*Bx;; Idgf3-OE* > ♂ *Ras*$^{V12}$

R-AD:
♀ *Bx* > ♂ *w*$^{1118}$
♀ *Bx* > ♂ *w*$^{1118}$*; Idgf3*$^{KD}$
♀ *Bx* > ♂ *w*$^{1118}$*, JNK*$^{DN}$
♀ *Bx;; αSpec-OE* > ♂ *w*$^{1118}$
♀ *Bx;; αSpec-OE* > ♂ *w*$^{1118}$*; Idgf3*$^{KD}$
♀ *Bx;; αSpec-OE* > ♂ *w*$^{1118}$*, JNK*$^{DN}$
♀ *Bx* > ♂ *Ras*$^{V12}$
♀ *Bx* > ♂ *w*$^{1118}$*; Idgf3*$^{KD}$*; Ras*$^{V12}$
♀ *Bx* > ♂ *w*$^{1118}$*, JNK*$^{DN}$*;; Ras*$^{V12}$
♀ *Bx;; αSpec-OE* > ♂ *Ras*$^{V12}$
♀ *Bx;; αSpec-OE* > ♂ *w*$^{1118}$*; Idgf3*$^{KD}$*; Ras*$^{V12}$
♀ *Bx;; αSpec-OE* > ♂ *w*$^{1118}$*, JNK*$^{DN}$*;; Ras*$^{V12}$
♀ *Bx, mCD8::RFP-OE* > ♂ *w*$^{1118}$*, JNK*$^{DN}$*;; Ras*$^{V12}$

**<u>Fig. 6</u>**

B-O:

♀ *Bx* > ♂ *w*$^{1118}$

♀Bx > ♂ w$^{1118}$; CH3L1-OE

♀*Bx* > ♂ *w*$^{1118}$*; CH3L2-OE*

♀*Bx* > ♂ *Ras*$^{V12}$

♀Bx > ♂ w$^{1118}$; CH3L1-OE; *Ras*$^{V12}$

♀*Bx* > ♂ *w*$^{1118}$*; CH3L2-OE; Ras*$^{V12}$

P:

♀ *Bx* > ♂ *w*$^{1118}$

♀Bx > ♂ w$^{1118}$; CH3L1-OE

♀*Bx* > ♂ *w*$^{1118}$*; CH3L2-OE*
